# Drinking-Water 4,4′-Methylenedianiline Establishes a Longitudinally Tractable Mouse Model of Cholestatic Liver Injury

**DOI:** 10.1101/2024.01.25.577198

**Authors:** Takumi Iwasaka, Katsuhisa Morita, Iori Azuma, Tomoka Nakagawa, Eri Nakashima, Tomomi Kamei, Yuki Kato, Hiroyuki Kusuhara, Tadahaya Mizuno

**Affiliations:** Graduate School of Pharmaceutical Sciences, the University of Tokyo, Bunkyo-ku, Tokyo, 113-0033, Japan; Laboratory for Drug Discovery and Development, SHIONOGI & Co.,Ltd. Toyonaka, Osaka, 561-0825, Japan; Graduate School of Pharmaceutical Sciences, the University of Tokyo, Bunkyo-ku, Tokyo, 113-0033, Japan, Electronic

**Keywords:** 4,4′-methylenedianiline, drinking-water model, cholestatic liver injury, toxicopathology, longitudinal profiling

## Abstract

Longitudinal studies of liver injury require models that can be maintained with simple administration, low procedure-related stress, and acceptable survival. Drinking-water thioacetamide (TAA-DW) is useful for this purpose, but it mainly produces centrilobular hepatocellular injury. Few drinking-water models are available for cholestatic liver injury. Here, we tested whether 4,4′-methylenedianiline (MDA), a known bile-duct-damaging hepatotoxicant, can be used to build a practical drinking-water model of cholestatic liver injury in mice. Male C57BL/6JJcl mice received MDA in drinking water for up to 28 days. A route- and administration-schedule-matched TAA-DW model was used as a reference. Drinking-water MDA (MDA-DW) caused sustained ALT elevation without observed mortality under the tested regimen. Compared with TAA-DW, MDA-DW produced stronger cholestatic biochemical features, including increased ALP, TBIL, and TCHO. Histology showed bile duct hyperplasia with degeneration, periportal/peribiliary fibrosis, and scattered focal hepatocellular necrosis. Whole-slide image analysis supported the distinct lesion distribution between MDA-DW and TAA-DW. Early time-resolved multimodal profiling showed model-specific responses: MDA-DW showed ALT elevation around Days 2–3, increased CD8-positive T cells on Day 8, and enrichment of fibrinolysis-related gene sets, whereas TAA-DW showed a sharp early injury peak and transient eosinophil increase. Impact Statement: This study expands the range of drinking-water liver injury models by providing a low-mortality MDA-DW model with cholestatic liver injury. The model offers a simple platform for time-resolved studies of liver injury progression, tissue remodeling, and model-specific inflammatory responses.

## Introduction

Liver injury is a major clinical and toxicological problem because repeated or persistent injury can lead to fibrosis, cirrhosis, liver failure, and cancer (Balakrishnan and Rehm 2024). Biliary and cholestatic injury is also an important form of liver injury, but it is difficult to study over time because many models require invasive procedures or specialized conditions (Mariotti et al. 2018). To understand how liver injury develops, adapts, and remodels over time, animal models should allow sampling at several time points with acceptable survival. This is especially important when the goal is to connect early biochemical, immune, and transcriptomic responses with later histological outcomes.

Chemical-induced liver injury models are widely used because the timing, route, and severity of injury can be controlled. Common models include carbon tetrachloride, acetaminophen, α-naphthylisothiocyanate, concanavalin A, and diet/chemical combinations (Tanaka et al. 2009; Jaeschke et al. 2012; Heymann et al. 2015; Scholten et al. 2015; Tsuchida et al. 2018). These models have been useful for studying hepatocellular injury, cholestatic injury, inflammation, fibrosis, and repair. However, many of them require repeated injections, gavage, special diets, or surgical procedures. These approaches are powerful, but they can add handling stress and technical variation, and they can make dense time-course studies difficult.

Drinking-water administration provides a simple way to deliver compounds continuously through the oral route. It reduces procedure-related stress and is well suited for time-resolved study designs. At present, however, the drinking-water model space for liver injury is limited. Thioacetamide administered in drinking water (TAA-DW) is useful and reproducible, but it mainly induces hepatocellular injury and fibrosis with a centrilobular pattern (Salguero Palacios et al. 2008; Wang et al. 2014). This creates a practical gap. Investigators can use TAA-DW to study centrilobular injury, but there are few simple drinking-water models that produce cholestatic liver injury.

We focused on 4,4′-methylenedianiline (MDA) because it offers an opportunity to convert a recognized toxicological phenotype into a practical drinking-water model. MDA is an aromatic amine with established hepatotoxicity and bile-duct liability. Human and animal studies have reported MDA-associated liver injury, biliary toxicity, and changes in biliary parameters (Kopelman et al. 1966; U.S. Department of Health and Human Services 1998; Dugas et al. 2001; Kwon et al. 2008). MDA has also been administered through drinking water in long-term carcinogenicity studies (Lamb et al. 1986). However, sustained MDA exposure through drinking water has not been systematically characterized as a mouse model for time-resolved liver injury studies. Thus, MDA drinking water (MDA-DW) may provide a practical way to extend drinking-water liver injury models toward sustained cholestatic liver injury with low observed mortality.

Here, we developed an MDA-DW model in mice and compared it with TAA-DW as a route- and administration-schedule-matched reference model. We first assessed liver injury, survival, and body-weight changes during 28 days of MDA-DW administration. We then characterized its biochemical and histological phenotype, including plasma liver tests, H&E and Sirius Red staining, and whole-slide image (WSI) analysis. Finally, we performed early time-resolved cross-sectional multimodal profiling using matched plasma and liver samples at each scheduled time point to compare the early response profiles of MDA-DW and TAA-DW. This study was designed to determine whether MDA-DW complements TAA-DW by expanding the drinking-water liver injury toolkit toward cholestatic liver injury and by providing a foundation for studying how biliary-pattern liver injury progresses and remodels over time in experimental systems.

## Methods

### Screening of candidate compounds for drinking water-induced liver injury model

Candidate compounds were screened using public toxicology databases, including CEBS, Open TG-GATEs, and DrugMatrix. As an initial broad filter, compounds associated with ALT values exceeding 200 U/L after any administration route or sampling time were extracted (Ganter et al. 2006; Waters et al. 2008; Igarashi et al. 2015). Candidate compounds were then refined by literature review using the following criteria: (a) evidence of liver injury after oral exposure, (b) water solubility compatible with drinking-water administration, defined as solubility of 1.0 g/L or greater, (c) practical availability and ease of handling, and (d) an expected injury profile distinct from TAA-DW, including biliary, periportal, or cholestatic features.

### Preparation of drug-induced liver injury models

Six-week-old male C57BL/6JJcl mice were purchased from CLEA Japan (Tokyo, Japan) and acclimatized for 3–5 days. Male mice were used as a practical choice for the initial characterization of the model, with the same sex used in the MDA and TAA groups. Mice were administered thioacetamide (TAA; T0817, Tokyo Chemical Industry Co., Japan; 300 mg/L) or 4,4′-methylenedianiline (MDA; M0220, Tokyo Chemical Industry Co., Japan; 750 mg/L) in drinking water to induce liver injury. Control mice received tap water. MDA free base was prepared at a nominal concentration of 750 mg/L in tap water by overnight stirring (10–20 h). This concentration was selected as a practical high concentration below the reported aqueous solubility of 1,000 mg/L at 25 °C (U.S. Department of Health and Human Services 1998), allowing preparation in water without a co-solvent. Drinking-water bottles were generally replaced every 4 days, with no interval longer than 7 days.

No a priori power calculation was performed. For this exploratory model-characterization study, we aimed to include at least four animals per group to obtain an initial assessment of between-animal variability. The numbers of animals included in each analysis are specified in the corresponding figure legends and tables.

Water intake was estimated from cage-level bottle weight loss, assuming a water density of 1 g/mL, and normalized by the number of animals and the measurement duration. A summary of estimated water intake over the initial observation intervals ending on Days 1–4 in Cohort 2 is presented in Table 2, with cumulative bottle-weight-loss estimates shown in Fig. S1. Mice were euthanized at each scheduled time point, and liver and blood samples were collected as described below. All animal procedures were approved by the Institutional Animal Care Committee of the Graduate School of Pharmaceutical Sciences, The University of Tokyo (approval number A2024P003-01), and were conducted in accordance with institutional guidelines and the Guiding Principles in the Use of Animals in Toxicology.

### Liver and blood sample collection

The first day of compound administration was defined as Day 0. Cohort 1 was used for biochemical and tissue characterization on Days 7, 14, and 28. Cohort 2 was sampled on Days 1, 2, 3, 4, and 8 for multimodal profiling. Both cohorts received drinking-water exposure from Day 0 until the scheduled endpoint. At each time point in Cohort 2, blood and liver samples were collected from the same animals. Separate groups of animals were used for terminal sampling at different time points.

In Cohort 2, under isoflurane anesthesia, blood was collected from the inferior vena cava into a 1.5 mL tube containing 1 µL heparin (Yoshindo Inc., Toyama, Japan). The superior vena cava was then clamped, and the portal vein was cut. The first liver perfusion was performed through the inferior vena cava with 10 mL Hanks’ Balanced Salt Solution containing 5 mM HEPES (H4034, Sigma-Aldrich) and 5 mM EDTA (345-01865, Fujifilm Wako Pure Chemical Corporation). The second perfusion was performed with 10 mL Hanks’ Balanced Salt Solution containing 5 mM HEPES. Before tissue harvest, 2 mL of gentleMACS dissociation enzyme solution (Miltenyi Biotec, Germany) was introduced into the liver through the inferior vena cava while the cut portal vein was clamped.

In Cohort 1, samples were collected on Days 7, 14, and 28. Blood was collected as described above. Liver perfusion was then performed through the inferior vena cava with 20 mL phosphate-buffered saline. For each animal, a portion of the outer left lateral lobe was collected for RNA isolation for RNA-seq or qPCR analysis. The remaining liver tissue was used for flow cytometry in Cohort 2 or histopathological analysis in Cohort 1. For longitudinal monitoring of blood biochemical values within the same individual, additional blood samples were collected from the tail vein where applicable. Details of data analysis are provided in the Supplementary Information.

### Analysis of blood specimens

Blood samples were centrifuged at 1,700 × g at 4 °C for 15 min to obtain plasma. Plasma levels of alanine aminotransferase (ALT), aspartate aminotransferase (AST), alkaline phosphatase (ALP), total bilirubin (TBIL), total cholesterol (TCHO), and triglycerides (TG) were measured using a DRI-CHEM NX500sV analyzer (Fujifilm Corporation, Japan). When a measured value exceeded the upper limit of the assay, the plasma sample was diluted with 100 mM HEPES containing 0.1% CHAPS (pH 7.4, room temperature) and re-measured. Reported values were corrected for the dilution factor.

### Analysis of liver specimens

Detailed procedures of RNA-seq analysis, histopathology and flow cytometry analysis, including subsequent data analysis, are provided in Supplementary Information.

### Quantitative PCR analysis

Total RNA was prepared using ISOGEN II (311-07361, Nippon Gene Co., Ltd., Tokyo, Japan). RNA was reverse-transcribed into cDNA using ReverTra Ace (FSQ-301, TOYOBO, Japan). Quantitative PCR was performed using THUNDERBIRD Next SYBR qPCR Mix (QPX-201, TOYOBO, Japan). Target gene expression was normalized to Gapdh. Primer sequences are listed in Table S1.

### Clustering Analysis

Dimensionality reduction was performed separately for the flow cytometry and transcriptome datasets obtained from liver samples on Days 1, 2, 3, 4, and 8 after administration. For each dataset, input features were reduced using principal component analysis (PCA), multidimensional scaling, t-distributed stochastic neighbor embedding, uniform manifold approximation and projection, spectral embedding, and locally linear embedding. The resulting low-dimensional representations were integrated using the META visualization method (Ma et al. 2023). The combined meta-distance matrix was visualized as a scatter plot, with each point color-coded by time, treatment condition, or ALT value. Transcriptome features were log-transformed before analysis. The code used for this analysis is available at GitHub (https://github.com/mizuno-group/mda-drinking-water-model). Analyses were performed using Python version 3.10.6 with open-source packages, including NumPy version 1.24.4, pandas version 2.0.3, SciPy version 1.11.1, scikit-learn version 1.1.2, statsmodels version 0.13.2, and umap-learn version 0.5.4.

### Gene set analysis

The Gene Ontology (GO) annotation file for mouse genes, m5.go.v2023.1.Mm.symbols.gmt, was downloaded from the MSigDB website and used for gene set analysis (Liberzon et al. 2011). Genes included in this annotation file were used as the analysis universe. Transcriptome data were log-transformed before analysis. Genes detected in fewer than 70% of samples, corresponding to fewer than 50 individuals, were removed. When zero-expression values were unevenly distributed across groups, this distribution was evaluated using one-way ANOVA applied to a binary zero-expression indicator, and genes showing group-biased zero expression were retained. Remaining missing values were imputed using KNNImputer in scikit-learn version 1.1.2.

The preprocessed RNA-seq data were analyzed using Generally Applicable Gene-set Enrichment (GAGE) to compare each treatment group with the corresponding control group at each time point (Luo et al. 2009). After applying GAGE, GO terms were summarized using the mean difference and adjusted P value using the Benjamini-Hochberg method for each model and time point. To prioritize GO terms that were consistently altered around the peak injury period, rank product statistics were calculated using mean difference values at 48 and 72 h (Eisinga et al. 2013).

## Data and code availability

The analysis code and tabular datasets used in this study are available at https://github.com/mizuno-group/mda-drinking-water-model. The version corresponding to this study is available under the release tag v1.0.0. Whole-slide images, derived histopathology features, and the MSigDB-derived gene-set reference are not included in the repository.

## Statistical analysis

Data are presented as mean ± standard deviation (SD) unless otherwise stated. Statistical and computational analyses were performed using Python version 3.10.6 with open-source packages. Transcriptome data were log-transformed before downstream analyses, and missing values were imputed using KNNImputer where applicable.

Comparisons between treatment groups and the corresponding controls at each time point were performed using two-sided Welch’s t-test without assuming equal variances unless otherwise specified. Plasma biochemical comparisons shown in Table 1 were performed using one-way ANOVA followed by Tukey’s HSD test. For immune-cell proportions in Fig. 4B and Fig. S4, groups were compared at each of the five time points for each of the eleven cell populations using the Kruskal–Wallis test. P values were adjusted across the 55 tests using the Benjamini–Hochberg procedure. Where the adjusted P value was below 0.05, Dunn’s test was used for the three pairwise comparisons, with Holm correction within each population–time-point set.

**Table 1:** Plasma biochemical values in Cohort 1 animals followed through Day 28.

| Day | group | BW | ALT | p-adj | AST | p-adj |
| --- | --- | --- | --- | --- | --- | --- |
| 7 | Control | 20.5±0.2 | 26.0±1.8 | - | 43.8±3.9 | - |
|  | TAA | 17.6±0.6 | 231.3±131.9 | * vs Control | 248.2±101.0 | ** vs Control |
|  | MDA | 19.0±1.1 | 326.3±65.5 | *** vs Control | 252.7±52.2 | ** vs Control |
| 14 | Control | 21.7±0.3 | 28.3±4.0 | - | 40.8±7.1 | - |
|  | TAA | 19.1±0.5 | 84.7±20.3 | ** vs Control | 88.3±19.7 | ** vs Control |
|  | MDA | 21.5±0.9 | 223.5±29.2 | *** vs Control, TAA | 178.2±23.2 | *** vs Control, TAA |
| 28 | Control | 23.7±0.2 | 24.3±0.5 | - | 32.5±1.3 | - |
|  | TAA | 20.7±0.8 | 76.3±18.6 | - | 61.7±12.5 | - |
|  | MDA | 21.1±0.5 | 135.3±75.8 | ** vs Control | 144.8±60.9 | ** vs Control, TAA |

Values are presented as mean ± SD for the same animals sampled on Days 7, 14, and 28 (n = 4 controls, n = 6 TAA-DW, and n = 6 MDA-DW). Statistical significance was assessed at each time point using one-way ANOVA followed by Tukey's HSD test. Asterisks indicate Tukey-adjusted P values: one asterisk, P < 0.05; two asterisks, P < 0.01; three asterisks, P < 0.001.

Correlations between plasma ALT levels and immune-cell proportions were evaluated using Pearson’s correlation coefficient. For WSI analysis, patch-level features extracted from hematoxylin and eosin (H&E)-stained liver sections were analyzed using a One-Class Support Vector Machine (One-Class SVM) trained on control-image patches with the expected anomaly fraction set to 0.05. Slide-level proportions of anomalous patches were compared against the reference value of 0.05 using a one-sample t-test for each treatment group and time point. Gene set enrichment analysis was performed using GAGE, and GO terms with a nominal P value < 0.05 were considered significantly enriched. Rank product statistics were calculated to identify GO terms consistently altered during the peak injury period. Unless otherwise stated, a P value < 0.05 was considered statistically significant.

## Results

### Development of MDA-DW as a Drinking-Water Model Distinct from TAA-DW

To develop a practical drinking-water model of liver injury that enables time-resolved investigation with minimal procedure-related stress, we screened candidate compounds from large-scale toxicology databases (Ganter et al. 2006; Waters et al. 2008; Igarashi et al. 2015). Compounds were prioritized based on established hepatotoxicity profiles, availability of chronic administration data, feasibility for oral or drinking-water exposure, potential for solubility and stability in aqueous media, and an expected injury profile distinct from TAA-DW. A literature survey, as detailed in the Methods section, guided the refinement process and led to the selection of MDA. MDA has been reported to exert pronounced hepatotoxicity and bile-duct liability, and it has previously been administered via drinking water in a carcinogenicity study in rats and mice (Kopelman et al. 1966; Lamb et al. 1986; Dugas et al. 2001; Kwon et al. 2008). Despite these findings, MDA has not been evaluated as a drinking water-based model compound for time-resolved experimental liver injury studies.

To examine its potential for inducing liver injury in mice, we administered MDA through drinking water over a 28-day period. In parallel, TAA, a widely used drinking-water hepatotoxicant model, was administered using the same general administration scheme to serve as a route- and schedule-matched reference model (Salguero Palacios et al. 2008; Wang et al. 2014). Plasma biochemical analyses were conducted on Days 7, 14, and 28 to assess liver injury. Both MDA- and TAA-treated groups showed significant elevations in ALT, a marker of hepatocellular injury (Fig. S2, Table 1, Table S2). Notably, the MDA-treated group consistently exhibited higher mean ALT levels than the TAA group across all time points, reaching statistical significance on Day 14. Under the tested regimen, no mortality was observed in the MDA-treated group.

To evaluate fibrotic responses, we performed qPCR for fibrosis-related genes. On Day 14, expression levels of *Col1a1* and *Acta2* were markedly increased in the MDA group compared to both the control and TAA groups (Fig. 1A, B), suggesting induction of fibrosis-related gene expression. Taken together, these results indicate that MDA administration via drinking water can reliably induce sustained liver injury with features distinct from TAA-DW.

**Fig. 1.**
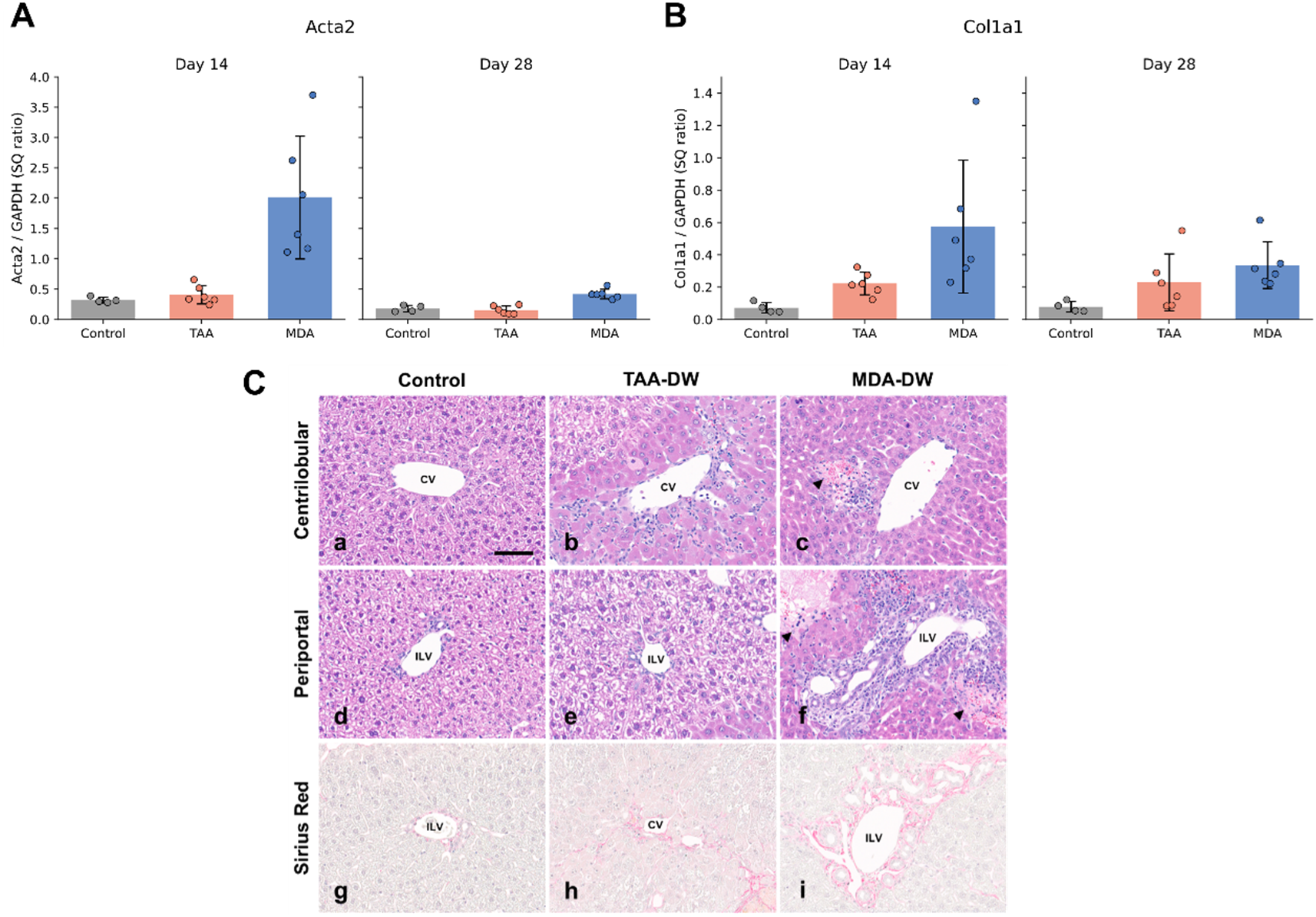
Characterization of MDA-DW and TAA-DW models in Cohort 1. (A, B) Quantitative PCR analysis of fibrosis-related genes on Days 14 and 28. Dots represent individual animals; bars and error bars show the mean ± SD. Sample sizes were n = 4 for controls and n = 6 for each treated group at each time point. (C) Representative H&E images of the centrilobular (a–c) and periportal (d–f) regions on Day 7 and Sirius Red images on Day 28 (g–i). Histopathological assessment included four animals per group and time point, except for TAA-DW on Day 14 (n = 3). Scale bar, 50 μm. Arrowheads indicate focal hepatocellular necrosis. CV, central vein; ILV, interlobular vein.

We also made a preliminary assessment of water intake using the available cage-level measurements from Cohort 2. Estimated intake was lower in the MDA-DW and TAA-DW groups than in controls over the initial observation intervals ending on Days 1–4 (Table 2; Fig. S1). At the Day 2 endpoint for TAA-DW and the Day 3 endpoint for MDA-DW, two treated cages were available for a descriptive assessment of between-cage variation; control cages were not replicated at these endpoints.

**Table 2:**
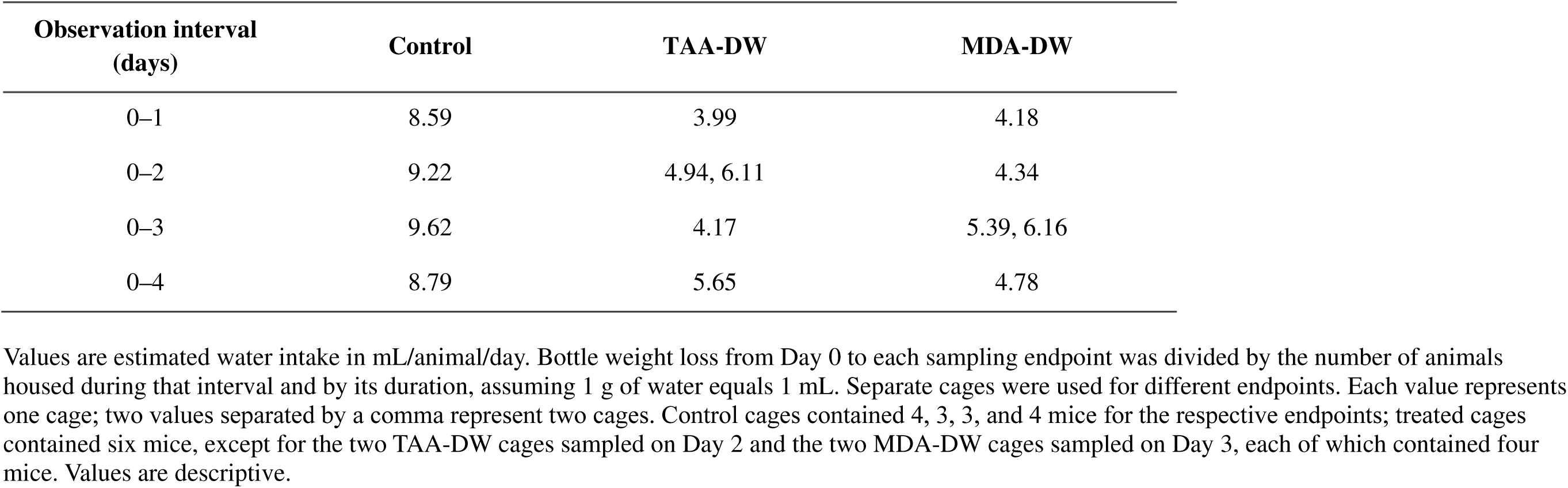
Estimated water intake over the initial observation intervals in Cohort 2.

| Observation interval<br>(days) | Control | TAA-DW | MDA-DW |
| --- | --- | --- | --- |
| 0–1 | 8.59 | 3.99 | 4.18 |
| 0–2 | 9.22 | 4.94, 6.11 | 4.34 |
| 0–3 | 9.62 | 4.17 | 5.39, 6.16 |
| 0–4 | 8.79 | 5.65 | 4.78 |

Values are estimated water intake in mL/animal/day. Bottle weight loss from Day 0 to each sampling endpoint was divided by the number of animals housed during that interval and by its duration, assuming 1 g of water equals 1 mL. Separate cages were used for different endpoints. Each value represents one cage; two values separated by a comma represent two cages. Control cages contained 4, 3, 3, and 4 mice for the respective endpoints; treated cages contained six mice, except for the two TAA-DW cages sampled on Day 2 and the two MDA-DW cages sampled on Day 3, each of which contained four mice. Values are descriptive.

### Histopathological Analysis

To further characterize the pathological features induced by MDA administration, liver tissues were collected on Days 7, 14, and 28 and subjected to H&E and Sirius Red staining (Fig. 1C). Histopathological scores summarizing lesion severity and frequency are presented in Table S3(a).

In the TAA-DW model, minimal to mild hepatocellular necrosis was predominantly observed in the centrilobular region, accompanied by mononuclear inflammatory cell infiltration and hepatocellular hypertrophy. These findings were present as early as Day 7 and remained detectable through Day 28. Oval cell hyperplasia, a regenerative response often associated with liver injury, was also evident in the centrilobular region, with a slight increase in both extent and frequency over time. These pathological features align with the known metabolic activation of TAA by CYP2E1, which is abundantly expressed in centrilobular hepatocytes and generates toxic intermediates that promote centrilobular injury (Ramaiah et al. 2001). Additionally, focal hemorrhagic necrosis was noted on the visceral surface of the left lateral lobe at Day 7, which subsequently evolved into regions rich in hemosiderin-laden macrophages by Days 14 and 28.

In contrast, the MDA-DW model demonstrated a markedly different lesion distribution. Hepatic injury was localized primarily to the periportal region, where mild to moderate bile duct hyperplasia with degenerative changes was consistently observed across all examined time points. Surrounding these ducts, inflammatory cell infiltration and periportal fibrosis were frequently detected. These findings are consistent with bile-duct-centered injury accompanied by a reactive ductular response and fibrotic remodeling (Dugas et al. 2001; Kwon et al. 2008). In addition, MDA-treated animals exhibited focal hepatocellular necrosis on Days 7, 14, and 28. Unlike the stereotyped zonal injury patterns seen with many hepatotoxins, these necrotic lesions lacked clear spatial consistency and appeared irregularly scattered throughout the parenchyma. On Days 14 and 28, focal macrophage clusters were observed at sites of necrosis, suggesting engagement of phagocytic clearance mechanisms. Hepatocellular atrophy was also present throughout the administration period.

To evaluate fibrosis with greater specificity, Sirius Red staining was employed. In the TAA-DW model, only minimal pericellular fibrosis was observed in centrilobular zones in one animal at Days 14 and 28. In contrast, the MDA-DW model exhibited peribiliary fibrosis as early as Day 7, and fibrosis remained detectable on Days 14 and 28. These observations were consistent with the fibrosis-related gene expression patterns described above and reinforce the distinction between the two models in terms of fibrosis distribution and severity. Detailed histopathological scoring is presented in (Table S3b–d).

### Unsupervised Analysis of Toxicopathological Profiles

As described above, irregularly scattered necrotic lesions were observed in the MDA-DW model, differing notably from the necrotic patterns typically associated with drug-induced liver injury. To characterize these toxicopathological features from an alternative perspective, we performed an unsupervised analysis using a computer-vision approach. First, each WSI was subdivided into smaller units, known as patches. We then extracted latent representations from each patch, converting them into numeric vectors suitable for analysis. An anomaly detection algorithm was subsequently applied to these patch vectors, enabling visualization of abnormal regions within each WSI in an unsupervised manner.

The TAA-DW model displayed extensive anomalous regions concentrated in a single contiguous area, whereas anomalous regions in the MDA-DW model were smaller and more diffusely scattered (Fig. 2A and B). Figure 2C and D show cropped images corresponding to the boxed regions in Fig. 2B. The cropped region from the TAA-DW model predominantly exhibited hemorrhagic necrosis, while irregularly scattered necrotic lesions were prominent in the MDA-DW model. Notably, the characteristic necrotic lesions in the MDA-DW model were relatively larger than typical cellular necrosis, as also evident at the WSI level (Fig. 2B). The proportion of patches classified as anomalous varied across groups and time points, with prominent values in the MDA-DW group on Days 14 and 28 (Fig. 2E).

**Fig. 2.**
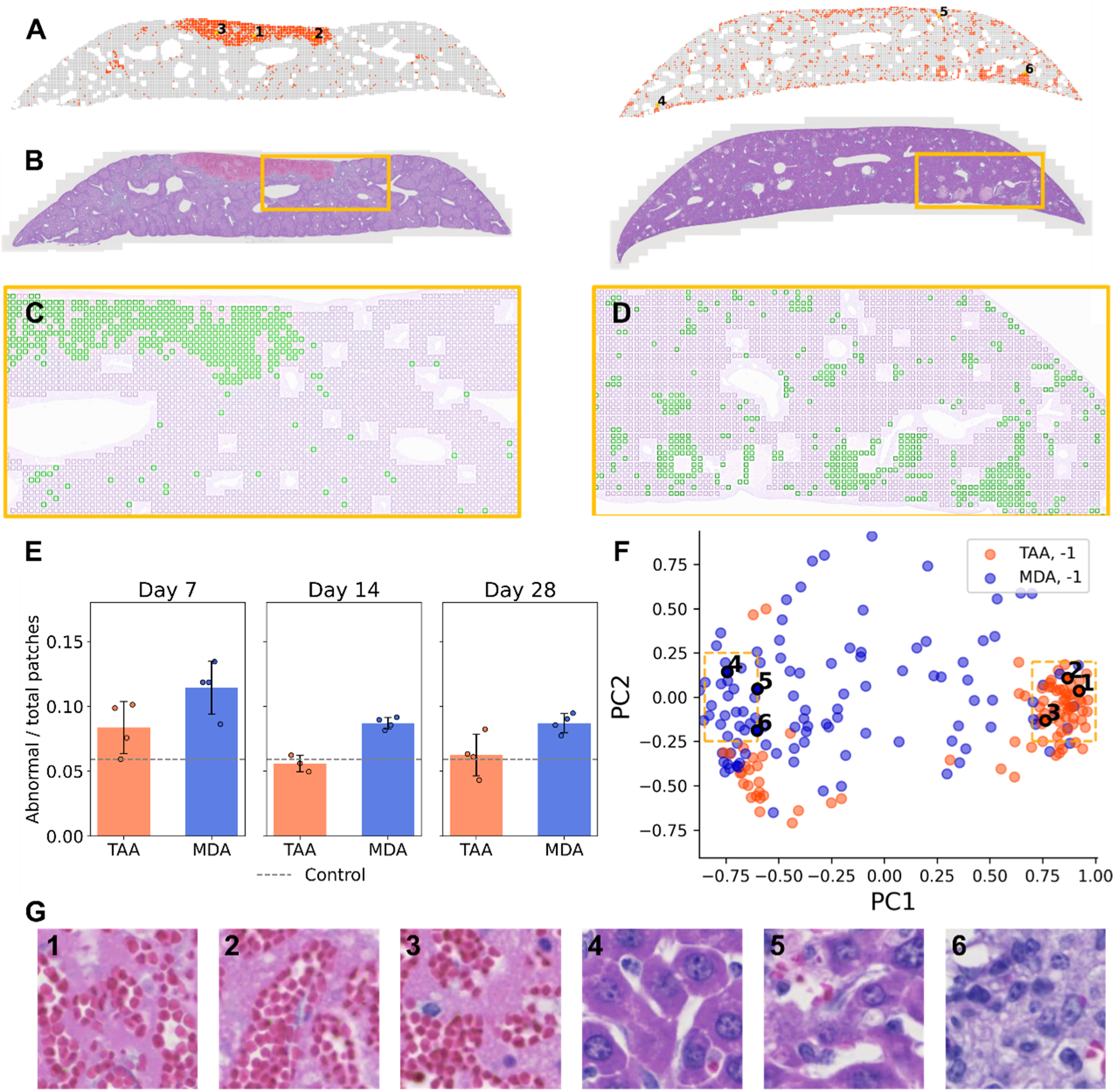
Unsupervised detection of abnormal regions in H&E-stained liver sections. (A) Patch-level anomaly detection results in whole-slide images. Red highlights indicate regions classified as anomalous. (B) Original H&E-stained whole-slide image. (C, D) Overlay images of the boxed regions in panel B. Green boxes indicate patches classified as anomalous. (E) Proportion of patches classified as abnormal, calculated for each slide as the number of abnormal patches divided by the total number of evaluated patches. Control values are shown as a reference; because control slides were included in model training, they were not treated as an independent group for inferential statistical testing. (F) PCA visualization of patch-level feature vectors and anomaly detection results on Day 7. Labels of 1 and −1 indicate normal and anomalous patches, respectively. (G) Representative cropped views of anomalous patches. Patch numbers correspond to those shown in panels A and F.

To investigate the heterogeneity of these lesions, we performed PCA on the patch vectors, revealing two distinct clusters corresponding to TAA-specific and MDA-specific lesions (Fig. 2F). Patch-level examination confirmed that TAA-specific clusters were characterized by extensive hemorrhagic necrosis, whereas MDA-specific clusters showed spatially dispersed focal hepatocellular necrosis without a clear zonal distribution (Fig. 2G).

Together, this analysis provides unsupervised quantitative support for the observation that the two models produce distinct pathological phenotypes.

### Early Time-Resolved Multimodal Profiling of MDA-DW and TAA-DW Models

Having established that the MDA-DW and TAA-DW models display clearly divergent pathological features over time, we next examined whether their early biological response profiles also differ. Our primary aim was to develop a drinking-water model suitable for time-resolved studies of liver injury. Therefore, we focused on the first 8 days after the start of administration, when biochemical injury, immune-cell changes, and transcriptional responses first emerge.

To compare these early responses, we designed a multimodal analysis focused on the early phase of each model (Fig. 3). Plasma biochemical markers were used to provide quantitative indicators of liver injury and systemic responses. Immune-cell profiling by flow cytometry was used to reveal shifts in innate and adaptive immune populations in the liver. Hepatic transcriptomic profiling was used to characterize molecular signatures associated with injury, inflammation, and tissue remodeling.

**Fig. 3.**
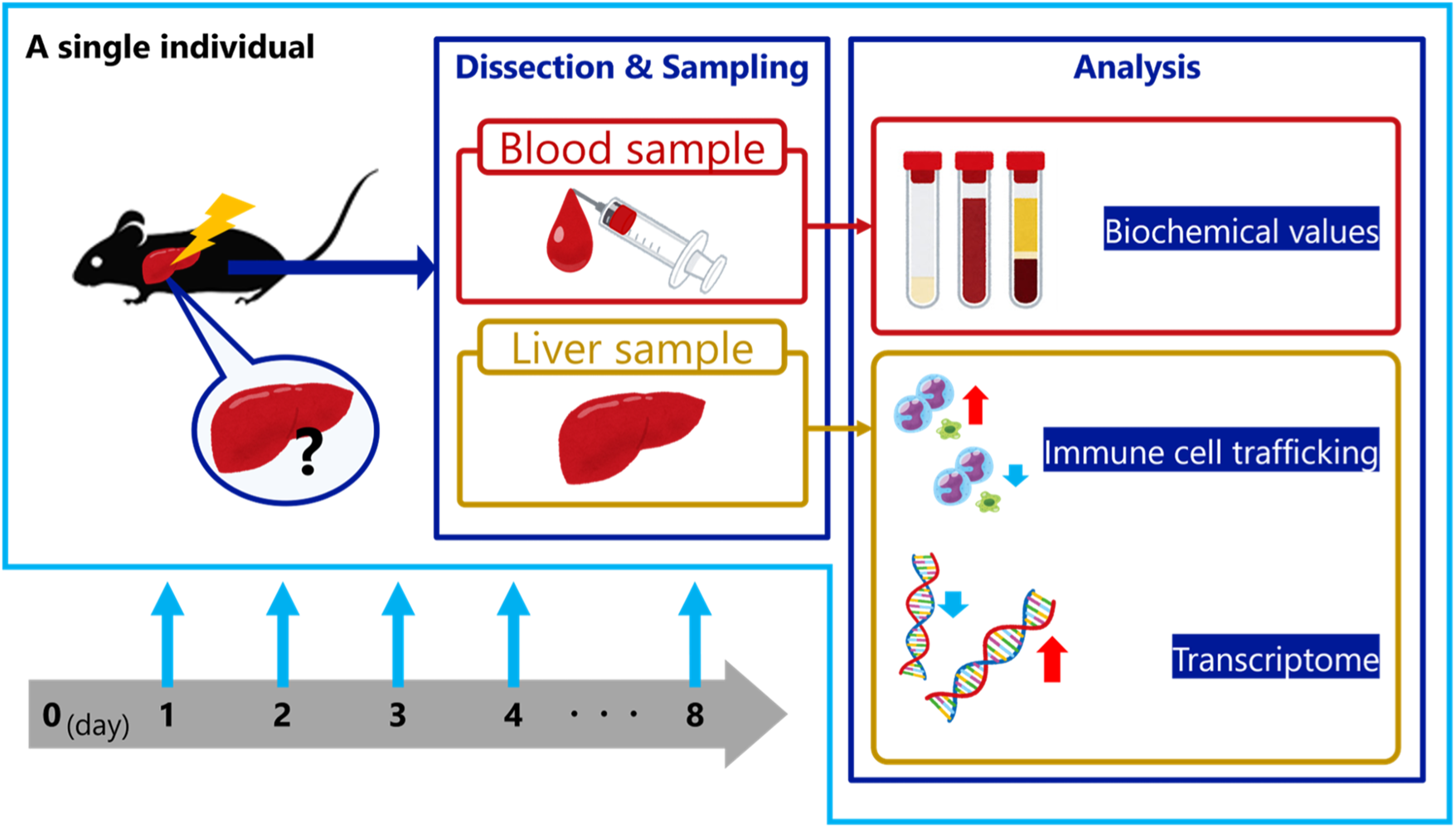
Multimodal sampling design for Cohort 2. At each scheduled time point, plasma and liver samples were collected from the same animals, enabling matched profiling of plasma biochemistry, hepatic immune-cell composition, and hepatic transcriptomes. Separate groups of animals were used for different time points; therefore, the design was time-resolved and cross-sectional rather than within-subject longitudinal profiling.

At each scheduled time point, plasma and liver samples were collected from the same animals, enabling matched multimodal profiling of plasma biochemistry, liver immune cells, and hepatic transcriptomes. Separate groups of animals were used for different time points; therefore, the analysis was time-resolved and cross-sectional rather than within-subject longitudinal profiling. This integrative approach allowed us to compare the emergence of model-specific biochemical, immunological, and molecular response patterns during the early phase of MDA-DW and TAA-DW injury (Fig. 3).

### Comparison of Plasma Biochemical Markers Between MDA-DW and TAA-DW Models

To assess systemic responses to liver injury, we first compared biochemical test results across the MDA-DW and TAA-DW models during the early administration phase. Both models showed elevated ALT levels indicative of hepatocellular damage, but the temporal dynamics and variability differed markedly (Table 3, Table S4). In the TAA-DW model, ALT levels peaked sharply on Day 2 with pronounced inter-individual variability, which then gradually declined over time. In contrast, ALT levels in MDA-DW were elevated around Days 2–3, with marked inter-individual variability on Day 3 (Table 3, Table S4). This suggests underlying heterogeneity in susceptibility to MDA-induced injury or differences in exposure kinetics.

**Table 3:** Plasma biochemical values in Cohort 2 on Days 1, 2, 3, 4, and 8.

| group | Day | BW | ALT | AST | ALP | TBIL | TCHO | HDL | TG |
| --- | --- | --- | --- | --- | --- | --- | --- | --- | --- |
| Control | 1 | 22.0±0.7 | 23.3±4.0 | 33.8±3.9 | 202.5±38.4 | 0.13±0.05 | **63.5±4.7** | 54.0±3.5 | 123.7±12.7 |
|  | 2 | 21.1±0.4 | 23.0±1.7 | 32.7±0.6 | **206.3±37.1** | 0.15 | 56.5 | 57.0 | 219.0 |
|  | 3 | 20.7±0.1 | **26.0±2.6** | 37.0±3.0 | 203.0±18.1 | 0.13±0.06 | 62.7±2.9 | 54.3±4.0 | 88.7±19.8 |
|  | 4 | 20.6±0.5 | 23.8±1.0 | 33.8±1.5 | 192.5±35.7 | 0.15±0.10 | 59.3±5.2 | 53.5±4.8 | 106.0±19.5 |
|  | 8 | 20.5±0.6 | 25.3±1.0 | **38.0±1.4** | 161.5±9.7 | **0.18±0.05** | 62.5±3.4 | 55.8±5.0 | 63.8±4.1 |
| MDA | 1 | 20.2±0.7 | 82.2±36.7 | 163.2±88.8 | 185.0±30.5 | 0.60±0.36 | 103.5±29.0 | 63.0±4.1 | **137.8±28.2** |
|  | 2 | 18.3±1.0 | **1403.3±336.8** | **1861.7±443.5** | 223.8±73.8 | 3.42±2.22 | 289.3±90.3 | 89.4±10.4 | 88.4±22.8 |
|  | 3 | 17.3±0.7 | 1388.8±1174.4 | 1205.0±1107.1 | 398.1±149.8 | **5.99±3.66** | 856.9±312.5 | **134.6±42.9** | 60.9±10.9 |
|  | 4 | 16.9±0.9 | 604.0±290.6 | 767.8±419.3 | 446.8±224.9 | 3.48±1.24 | **1112.3±630.9** | 80.5±29.2 | 48.8±14.2 |
|  | 8 | 19.9±2.4 | 393.3±92.6 | 261.3±62.2 | **1095.0±333.9** | 0.97±0.30 | 710.5±734.7 | 116.0±46.0 | 90.3±48.2 |
| TAA | 1 | 20.9±0.7 | 62.2±27.9 | 70.8±24.7 | 287.5±42.5 | 0.15±0.05 | 76.8±7.3 | 57.6±2.9 | **83.0±26.5** |
|  | 2 | 18.8±1.7 | **21197.5±20513.4** | **16836.0±16630.7** | 308.4±56.3 | 0.49±0.27 | 63.5±18.8 | 52.0±14.8 | 55.3±32.4 |
|  | 3 | 18.8±0.9 | 4463.3±1120.9 | 2523.3±1343.4 | 264.8±30.3 | 0.48±0.18 | 78.0±27.2 | 43.5±18.5 | 24.7±9.8 |
|  | 4 | 17.0±0.4 | 2862.7±1419.2 | 2733.0±1632.3 | **338.8±155.1** | **0.75±0.26** | 71.0±22.0 | 38.7±14.6 | 34.3±15.7 |
|  | 8 | 18.2±1.9 | 231.0±299.2 | 299.8±222.8 | 164.5±43.3 | 0.43±0.15 | **90.3±21.8** | **61.8±9.7** | 76.0±28.8 |

Values are presented as mean ± SD. For parameters with only one or two samples (n = 1 or 2), values represent the mean without SD. Bold values indicate peak measurements observed during the treatment period under each condition.

Despite continued compound administration, both models exhibited partial attenuation of ALT elevation by Day 8. Importantly, the MDA-DW group also exhibited significant elevations in ALP, TBIL, and TCHO, biochemical markers associated with cholestatic liver dysfunction, which were less pronounced in the TAA-DW group (Table 3, Fig. S3). These findings are consistent with previous reports describing cholestatic features in MDA-induced toxicity following single-dose oral administration (Kwon et al. 2008). Collectively, the MDA-DW and TAA-DW models demonstrate distinct early biochemical profiles, with stronger cholestatic biochemical features in MDA-DW.

### Comparison of Immune Cell Profiles in MDA-DW and TAA-DW Models

Given the central role of immune responses in liver injury and tissue remodeling (Hatada et al. 2005; Saijou et al. 2018; Hossain and Kubes 2019; Yang and Tonnesseen 2019; Liu et al. 2021), we next investigated immune-cell dynamics in the two models using flow cytometry. To explore global changes over time, we applied META visualization, an ensemble method integrating multiple dimensionality reduction techniques, to cytometry data, enabling identification of broad immunological trends (Ma et al. 2023) (Fig. 4A). Although treatment and control groups were indistinguishable in the early phase, temporal divergence emerged with continued exposure.

**Fig. 4.**
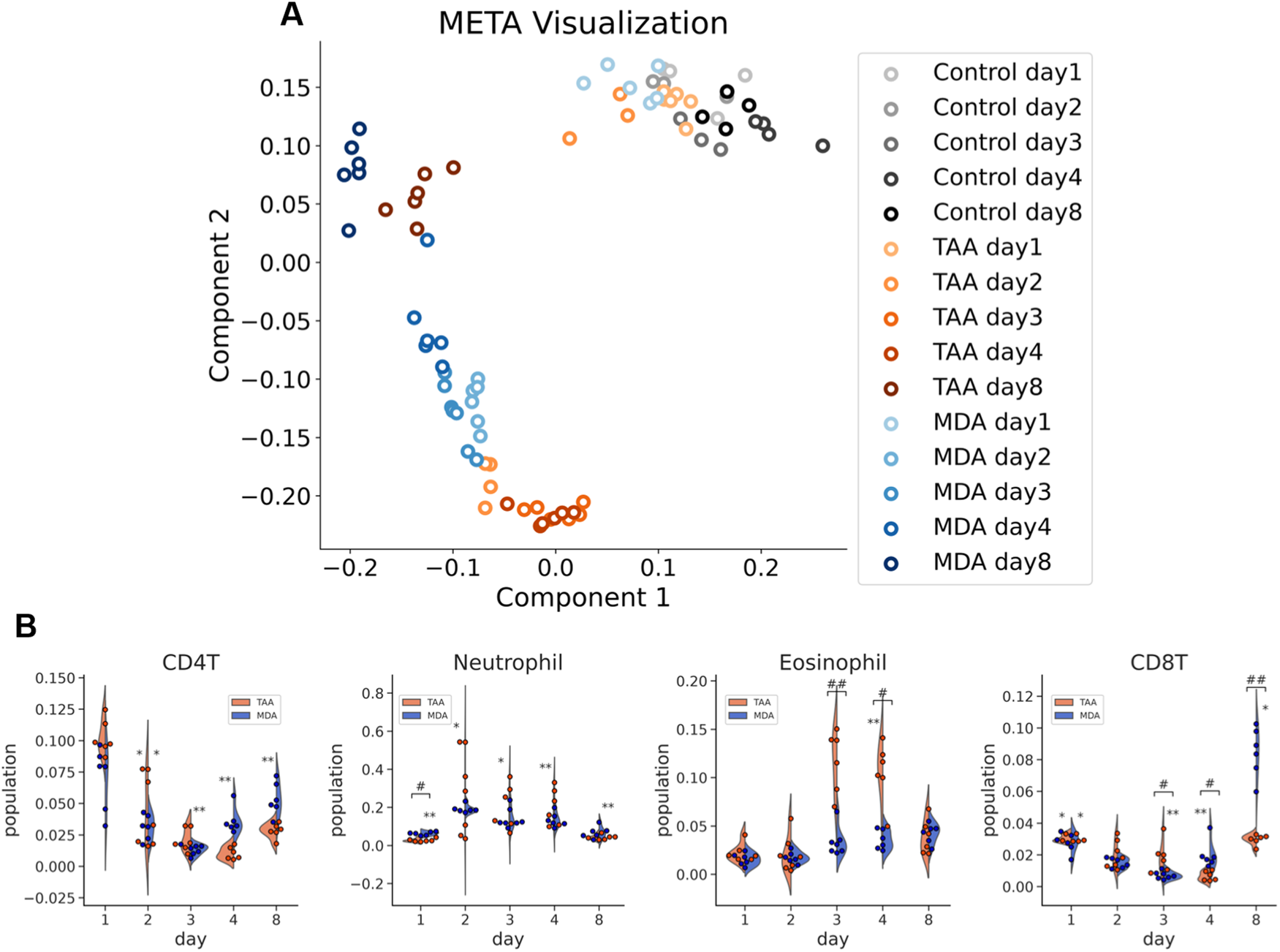
Comparison of hepatic immune-cell profiles between MDA-DW and TAA-DW models. (A) Flow cytometry-derived immune-cell profiles were embedded in two dimensions using META visualization. (B) Violin plots show the proportions of selected hepatic immune-cell populations at each time point. Dots represent individual animals. The x-axis is categorical and is not scaled according to elapsed time. Statistical analysis used Kruskal–Wallis tests with Benjamini–Hochberg correction across all 55 population–time-point tests in Fig. 4B and Fig. S4, followed, where the adjusted P value was below 0.05, by Dunn’s pairwise tests with Holm correction within each population–time-point set. Asterisks indicate differences from the concurrent control group, which is included in the statistical analysis but not displayed. Hash marks indicate differences between MDA-DW and TAA-DW. One, two, and three symbols indicate Holm-adjusted P values < 0.05, < 0.01, and < 0.001, respectively.

Zoomed-in analyses revealed shared and model-specific immune-cell behaviors. Both models showed a reduction in lymphocyte populations and an increase in neutrophil proportions around the times of peak ALT elevation (Fig. 4B). Kupffer cell proportions were lower than in controls in TAA-DW on Days 2–4 and in MDA-DW on Day 8 (Fig. S4).

In addition to shared responses, clear model-specific differences were identified. Eosinophil proportions were higher in TAA-DW than in controls on Day 4 and higher than in MDA-DW on Days 3 and 4. Conversely, CD8-positive T-cell proportions were higher in MDA-DW than in both controls and TAA-DW on Day 8. These findings indicate that while both models activate shared inflammatory responses, they diverge in the recruitment or involvement of specific immune subsets, potentially reflecting differences in the spatial context and nature of tissue injury.

### Comparison of Hepatic Transcriptome Profiles in MDA-DW and TAA-DW Models

To examine molecular-level differences in the hepatic response to injury, we performed RNA sequencing on liver tissues from both models. Gene expression data were visualized using META visualization (Fig. 5A, B; see also Fig. S5). Samples collected shortly after compound administration clustered near control animals but diverged progressively over time. By Day 8, individuals with lower ALT levels tended to shift back toward the control cluster, whereas those with sustained liver injury remained distant. This pattern was observed regardless of the model, suggesting a close association between functional injury and transcriptional disruption.

**Fig. 5.**
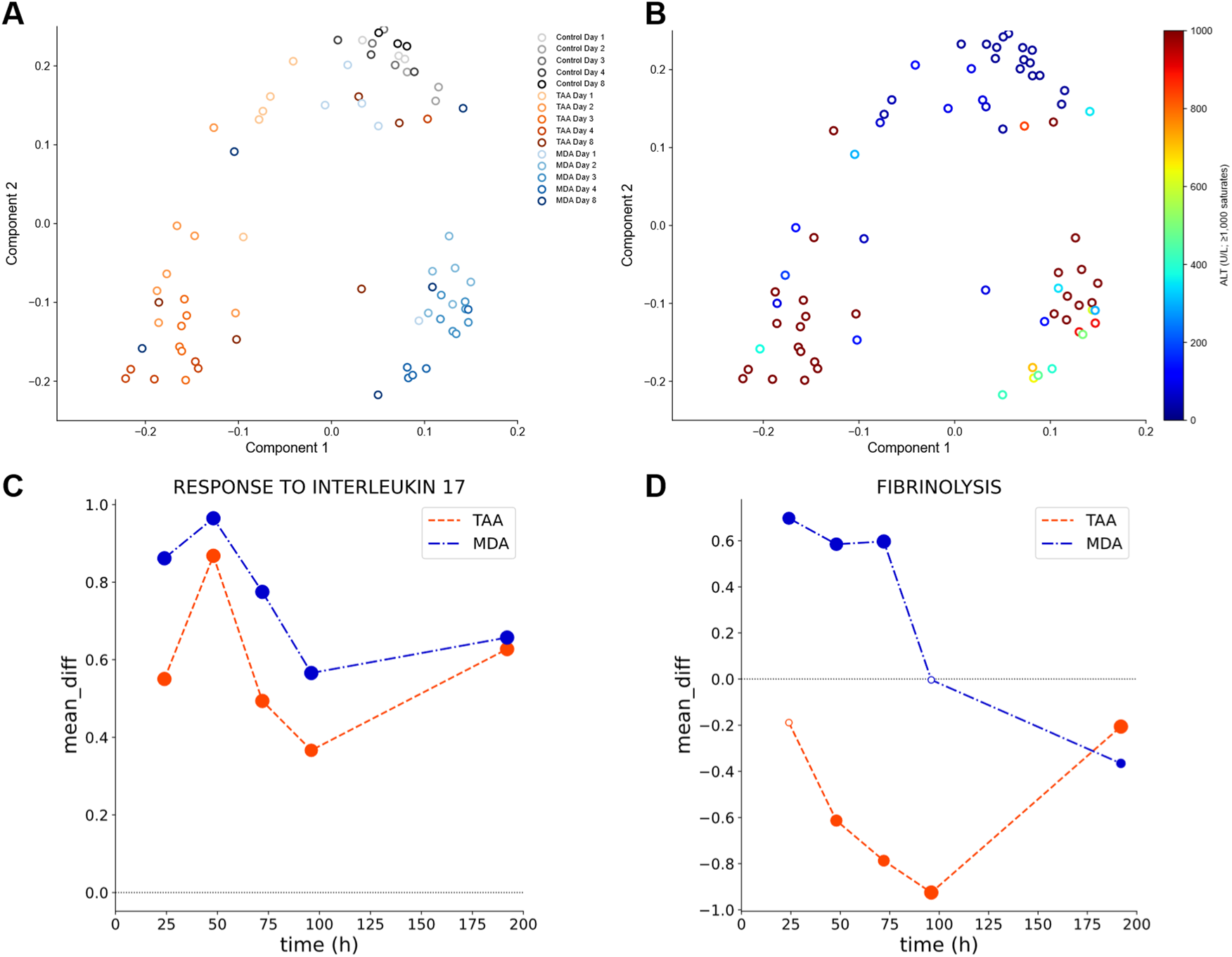
Comparison of hepatic transcriptome profiles between MDA-DW and TAA-DW models. (A, B) RNA-seq profiles were embedded in two dimensions using META visualization and colored by treatment/time point (A) or plasma ALT level (B). In (B), ALT values ≥1,000 U/L are displayed at the upper end of the color scale. In (C), circle size represents −log(P) from GAGE, whereas filled symbols indicate GO terms with Benjamini–Hochberg–adjusted P < 0.05 within each treatment–time condition. Circle size corresponds to the −log(P value) calculated by GAGE.

The transcriptomic divergence between models increased over time. On Day 3, distinct clusters were observed for MDA-DW and TAA-DW animals, underscoring model-specific gene expression responses to each compound. When visualized alongside ALT levels, animals with lower enzyme levels consistently resided closer to control-like transcriptomic states (Fig. 5B), suggesting that ALT broadly tracked the overall degree of molecular disruption.

To identify key biological processes associated with these transcriptomic shifts, we conducted gene set analysis using the GAGE framework, which accommodates unbalanced sample sizes (Luo et al. 2009). In the MDA-DW model, IL-17-related and monocyte-associated GO terms were significantly enriched during the acute phase, especially on Days 2 and 3, consistent with our immune profiling results (Fig. 5C, Table S5). The TAA-DW model exhibited a similar enrichment trend, suggesting that some inflammatory programs were shared across injury types (Table S6).

However, notable divergences emerged in matrix-remodeling-related pathways. At the time points shown in Fig. 5C, fibrinolysis-related GO responses differed between the models, with positive changes in MDA-DW and negative changes in TAA-DW (Table S6). At the transcriptome level, these findings suggest that matrix-remodeling-related molecular signatures differ depending on the nature of liver injury. These molecular differences further reinforce the histological and immunological distinctions observed between the models.

We also examined selected genes related to bile acid synthesis and hepatobiliary transport in the existing RNA-seq dataset (Fig. S6). In MDA-DW, *Cyp7a1* expression was lower on Day 1, whereas *Abcc3*, *Slc51b*, and *Abcb4* showed higher expression than controls. These observations indicate changes in bile acid–related gene expression during the early response period.

### Integrated Characterization of Complementary Drinking-Water Liver Injury Models

Together, these results reveal that the MDA-DW and TAA-DW models represent complementary drinking-water liver injury models, each characterized by distinct biochemical, histological, immune-cell, and transcriptomic responses (Fig. 6). While both models induce liver injury through drinking-water administration, their early-phase trajectories diverge across multiple biological layers: plasma biochemical markers, immune-cell dynamics, and hepatic transcriptome signatures.

**Fig. 6.**
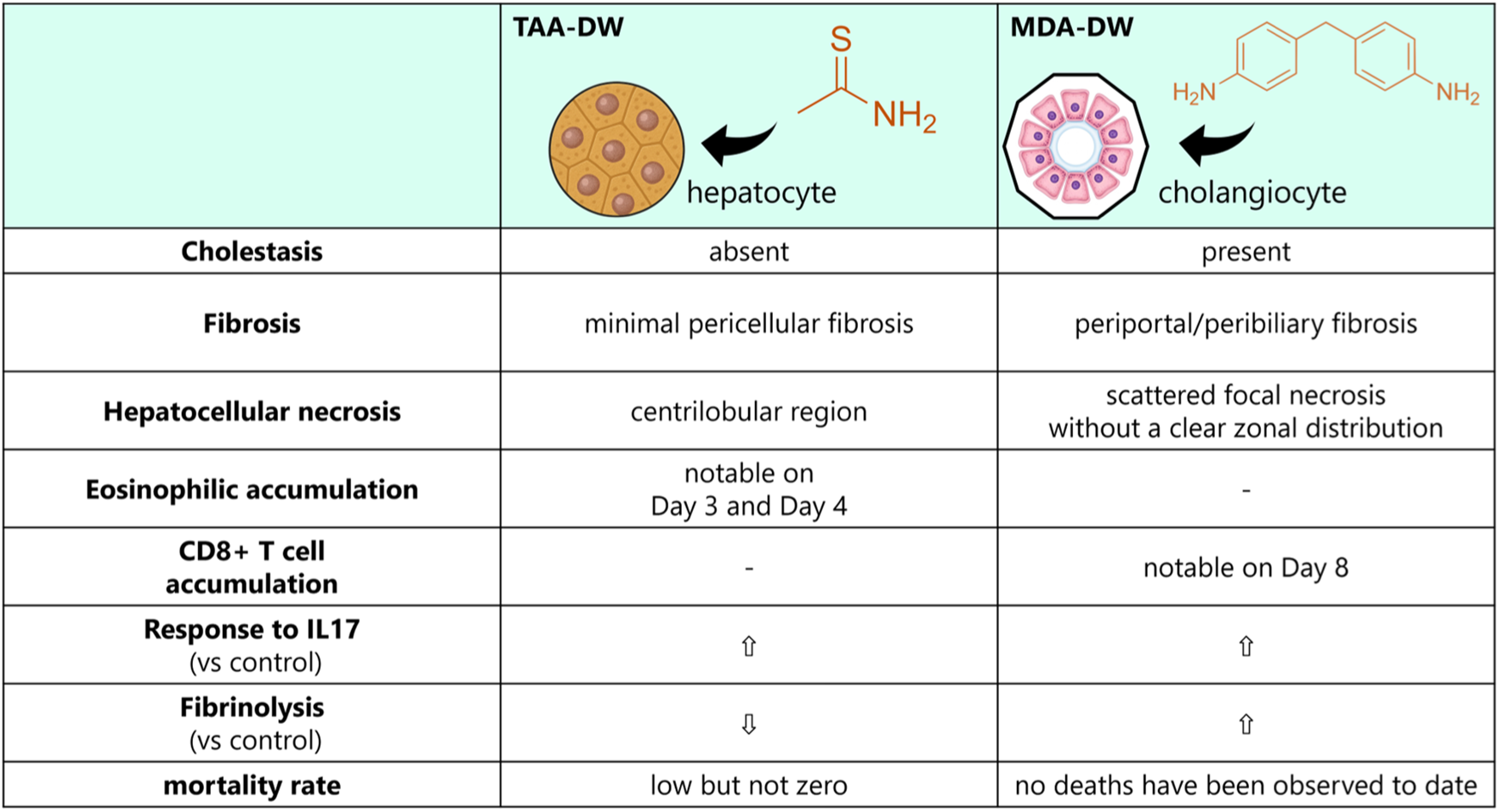
Integrated comparison of complementary MDA-DW and TAA-DW liver injury models. Summary of the major biochemical, histological, immune-cell, and transcriptomic features observed in the two drinking-water models. Upward and downward arrows indicate the direction of changes in gene-set expression relative to controls during the early response period.

TAA-DW was characterized by a sharp early ALT peak, centrilobular hepatocellular injury, and transient eosinophil enrichment. In contrast, MDA-DW was characterized by sustained liver injury under the tested regimen, cholestatic biochemical features, bile duct-centered histopathology, spatially dispersed focal necrosis, increased CD8-positive T cells on Day 8, and upregulation of the fibrinolysis-related gene set during Days 1–3. These multi-layered differences align with the divergent phenotypes observed in the two models, including the zonal distribution of necrosis, fibrosis localization, and cholestatic features.

A multi-layered comparison of these models highlights their utility as complementary platforms for studying diverse patterns of liver injury. In particular, MDA-DW extends the available drinking-water liver injury toolkit toward cholestatic liver injury, providing a practical foundation for time-resolved studies of liver injury progression and tissue remodeling in experimental systems.

## Discussion

In this study, we developed MDA-DW as a simple drinking-water mouse model that complements the existing TAA-DW model. The main contribution of this study is the conversion of the known bile-duct liability of MDA into a practical model platform for time-resolved studies of cholestatic liver injury. Under the tested regimen, MDA-DW induced sustained liver injury with low observed mortality, cholestatic biochemical features, bile duct-centered histopathology, spatially dispersed focal necrosis, and model-specific immune and transcriptomic responses. These findings show that MDA-DW extends the drinking-water liver injury toolkit beyond the centrilobular hepatocellular injury pattern represented by TAA-DW.

A key feature of MDA-DW was its distinct histopathological pattern. TAA-DW mainly produced centrilobular hepatocellular injury, consistent with the known metabolic activation of TAA in centrilobular hepatocytes (Ramaiah et al. 2001). In contrast, MDA-DW produced bile duct hyperplasia with degenerative changes, periportal inflammatory cell infiltration, periportal/peribiliary fibrosis, and focal hepatocellular necrosis without a clear zonal distribution. This pattern is consistent with previous reports describing MDA-associated bile duct injury and altered biliary parameters (Dugas et al. 2001; Kwon et al. 2008). The value of the present study is that sustained drinking-water MDA exposure provides a tractable mouse model with a cholestatic liver injury phenotype that is distinct from TAA-DW.

The spatial distribution of injury further supports the distinction between the two models. In TAA-DW, abnormal regions detected by WSI analysis were concentrated in large contiguous areas, mainly corresponding to hemorrhagic or centrilobular necrotic lesions. In MDA-DW, abnormal regions were smaller and more diffusely distributed, corresponding to scattered focal injury regions. This unsupervised analysis provided quantitative support for the pathologist-based observation that the two drinking-water models differ in lesion topology. Such spatial information is important because lesion location can influence immune-cell recruitment, tissue remodeling, and the interpretation of plasma biochemical markers.

MDA-DW also showed biochemical and histological features that were more biliary or cholestatic in nature than those of TAA-DW. MDA-DW increased ALP, TBIL, and TCHO, together with bile duct changes and periportal/peribiliary fibrosis. These features support the use of MDA-DW as a model exhibiting cholestatic liver injury phenotypes. MDA-DW also showed a higher proportion of hepatic CD8-positive T cells on Day 8. CD8-positive cytotoxic T cells have been implicated in immune-mediated cholangiopathies, including Primary Biliary Cholangitis and autoimmune cholangitis models, where they are associated with portal inflammation, bile duct injury, and biliary epithelial cell damage (Kita et al. 2002; Li et al. 2019; Zhu et al. 2024). Thus, MDA-DW may capture selected biliary, cholestatic, and immune features that are relevant to immune-mediated cholangiopathy research. At the same time, MDA-DW should not be considered a complete model of a specific human cholangiopathy. Direct comparisons with established cholestatic, toxic, and autoimmune cholangiopathy models, such as BDL, ANIT, DDC, and genetic or immune-mediated models, will be needed to define which disease-relevant components are shared with MDA-DW (Mariotti et al. 2018).

Biliary exposure to MDA metabolites is a possible contributor to cholangiocyte injury, as studies in rats have linked MDA toxicity with its disposition and biliary excretion (Dugas et al. 2001). The additional RNA-seq analysis showed changes in bile acid–related transcripts, including a transient decrease in *Cyp7a1* and increases in *Abcc3*, *Slc51b*, and *Abcb4* (Fig. S6). These changes may reflect a response involving farnesoid X receptor (FXR)-regulated pathways (Zollner et al. 2006). Their functional significance and relationship to the histological findings in Cohort 1 remain to be established. Measurements of bile acids, bile flow, transporter function, and MDA metabolites would help clarify the mechanism.

Multimodal profiling in Cohort 2 showed distinct response patterns in MDA-DW and TAA-DW during Days 1–8. In plasma biochemistry, TAA-DW showed a sharp early ALT peak, whereas MDA-DW showed ALT elevation around Days 2–3 and stronger cholestatic biochemical changes. In immune-cell profiling, both models shared inflammatory features, including increased neutrophil proportions associated with ALT elevation and changes in Kupffer cell proportions. At the same time, model-specific immune changes were observed: eosinophils increased transiently in TAA-DW, whereas CD8-positive T cells increased on Day 8 in MDA-DW. These differences suggest that the two models are not only histologically distinct but also differ in their early inflammatory response patterns.

Transcriptome profiling further supported this model-specific divergence. IL-17-related and monocyte-associated gene sets were enriched during the early phase, indicating that inflammatory programs were activated in both models. In contrast, fibrinolysis-related GO terms showed opposite directions of regulation: upregulated in MDA-DW and downregulated in TAA-DW. Fibrin deposition and fibrinolysis have been linked to liver injury, ductular reactions, and cholestatic injury models, including ANIT-based models (Luyendyk et al. 2009; Kopec and Luyendyk 2014; Kopec et al. 2017). Therefore, the fibrinolysis-related transcriptomic signature observed in MDA-DW may reflect a matrix-remodeling response associated with biliary-pattern injury. However, this interpretation remains provisional because fibrin deposition was not directly examined histologically in the present study.

An important practical advantage of MDA-DW is its suitability for time-resolved experimental designs. Many liver injury models require repeated injections, gavage, surgery, special diets, or genetic manipulation, which can add procedure-related stress and limit dense sampling schedules. Drinking-water administration provides continuous oral exposure with minimal handling. In the present study, plasma and liver samples were collected from the same animals at each scheduled early time point, allowing matched multimodal profiling of plasma biochemistry, liver immune cells, and hepatic transcriptomes. Separate groups of animals were used across time points, so the design was time-resolved and cross-sectional rather than within-subject longitudinal. This distinction is important, but the design still provides a practical framework for connecting early biochemical, immune, and molecular responses with later histological outcomes.

This study has several limitations. We did not directly benchmark MDA-DW against established cholestatic or cholangiopathy models, such as BDL, ANIT, DDC, or genetic/immune-mediated models; future comparisons will clarify which disease-relevant features are shared or unique. MDA concentration and stability in drinking water were not analytically verified, and water-intake estimates were not corrected for evaporation or leakage. Because compound intake depends on both water concentration and consumption, actual MDA intake remains uncertain. Food intake was not measured, and both water and food intake should be monitored when applying this regimen. MDA toxicokinetics, bile acid profiles, transporter protein expression and function, and fibrin deposition were not directly assessed. The study used male mice under a single exposure regimen without a withdrawal arm, and small group sizes limit the precision of the estimates. Given the greater susceptibility reported in female rats (Dugas et al. 2001), evaluation in both sexes is needed. The observed biliary, immune, and transcriptomic features provide endpoints for future comparisons and targeted mechanistic studies.

In conclusion, MDA-DW provides a simple drinking-water mouse model with low observed mortality under the tested regimen and a cholestatic liver injury phenotype distinct from TAA-DW. By integrating plasma biochemistry, histopathology, WSI analysis, liver immune-cell profiling, and transcriptomics, this study defines MDA-DW and TAA-DW as complementary drinking-water liver injury models. MDA-DW extends the available model toolkit toward biliary-pattern injury and provides a practical foundation for time-resolved studies of liver injury progression and tissue remodeling in experimental systems.

## Acknowledgements

We thank all those who contributed to the construction of the data sets employed in the present study such as CEBS, Open TG-GATEs, and DrugMatrix. We also thank Dr. Toru Komatsu (the University of Tokyo) for helpful discussion.

## Author Contributions

TI: Data curation, Formal analysis, Investigation, Writing – Original draft, Visualization.

KM: Investigation, Data curation

IA: Investigation, Data curation

TN: Investigation

EN: Investigation

TK: Formal analysis

YK: Formal analysis, Writing – Original draft, Writing – Review and editing

HK: Writing – Review and editing.

TM: Conceptualization, Resources, Supervision, Project administration, Writing – Original draft, Writing – Review and editing, Funding acquisition.

## Financial Support and Sponsorship

This work was supported by JSPS KAKENHI Grant-in-Aid for Scientific Research (C) (Grant Number 21K06663), JSPS KAKENHI Grant Number JP22H04925 (PAGS), Takeda Science Foundation, and Mochida Memorial Foundation for Medical and Pharmaceutical Research.

## Data Deposit and Linking

The raw RNA-seq sequencing data generated in this study have been deposited in the DNA Data Bank of Japan (DDBJ) under BioProject accession number PRJDB14915. The sequencing data are associated with DDBJ Sequence Read Archive (DRA) accession DRA016140.

## Statement

During the preparation of this work the author(s) used Paperpal for English editing. After using this service, the author(s) reviewed and edited the content as needed and take(s) full responsibility for the content of the publication.

## Conflicts of Interest

TK and YK are employees of Shionogi & Co., Ltd., which partially funded this study. All other authors declare no competing interests.

## List of Abbreviations

MDA: 4,4’-methylenedianiline;
TAA: thioacetamide;
ALT: alanine aminotransferase;
AST: aspartate aminotransferase;
GO: Gene Ontology;
GAGE: Generally Applicable Gene-set Enrichment;
ANIT: α-naphthylisothiocyanate;
HSCs: hepatic stellate cells;
PFs: portal fibroblasts

## Supplementary Information

### Supplementary Methods

#### RNA-seq analysis

Poly(A)-tailed mRNA was purified from total RNA prepared with ISOGEN II (311-07361, Nippon Gene, Japan) using the NEBNext Poly(A) mRNA Magnetic Isolation Module. RNA-seq libraries were prepared using the NEBNext Ultra II Directional RNA Library Prep Kit for Illumina. Libraries were sequenced using the NovaSeq 6000 S1 Reagent Kit v1.5 (200 cycles).

Quality control of reads was performed using PRINSEQ++ version 1.2.4 with the following parameters: trim_left=5, trim_tail_right=5, trim_qual_right=30, ns_max_n=20, and min_len=30. Transcript abundance was quantified using Salmon version 1.6.0 with GENCODE vM28 transcript annotations and the GRCm39 primary assembly genome (1). Salmon was run with validateMappings, gcBias, and seqBias options using a decoy-aware index (2). Transcript-level quantification files, quant.sf, were imported into R version 4.2.3 using tximport in Bioconductor. Scaled Transcript per million values were used for downstream analysis.

For the supplementary analysis, expression profiles of selected genes related to bile acid synthesis, hepatobiliary transport, and biliary epithelial cells were extracted from the existing liver RNA-seq dataset and displayed by treatment and sampling time. This analysis was descriptive.

#### Histopathology

The left lateral lobe of the liver was excised and collected for histological analysis into a 5 mL tube containing 4 mL of 10% neutral-buffered formalin. Samples were fixed for 24 hours, washed with PBS for 2 hours, and subsequently stored in 70% ethanol. Tissue processing, sectioning, and staining with hematoxylin and eosin (H&E) or Sirius Red were outsourced to Genostaff Inc.

Tissue sections were prepared by longitudinally slicing the left lateral lobe. Fixed liver tissues were subjected to a standard dehydration and paraffin embedding protocol. Samples were first immersed in 70% ethanol at 4 °C for 3 days, followed by sequential incubations in 80%, 90%, and 100% ethanol for 1 hour each at room temperature (RT).

Subsequent processing was performed using an automated tissue processor (CT-Pro20, Genostaff Inc.). Tissues underwent three consecutive immersions in 100% ethanol (1.5 hours each at RT), followed by incubation in a 1:1 mixture of 100% ethanol and G-Nox (xylene substitute, Genostaff Inc.) for 1.5 hours at RT. Samples were then immersed sequentially in three changes of G-Nox (1.5 hours each at RT), and finally in three changes of paraffin wax at 65 °C for 1.5 hours each.

Paraffin blocks were sectioned at a thickness of 4 µm and used for both H&E and Sirius Red staining.

For H&E staining, tissue sections were deparaffinized with xylene, rehydrated through a graded ethanol series, and rinsed in running tap water for 5 minutes, followed by distilled water. Sections were stained with hematoxylin (Sigma) for 3 minutes, rinsed in running tap water for 20 minutes, stained with eosin alcohol (Sigma) for 3 minutes, and then mounted with Malinol (#2040, Muto).

For Sirius Red staining, deparaffinized tissue sections were stained in Iron Hematoxylin Working Solution (Weigert’s Iron Hematoxylin #4034, Muto Pure Chemicals) for 3 minutes at RT, treated with hydrochloric acid ethanol solution for 3 minutes, and rinsed in running tap water followed by distilled water. After treatment with 5% phosphomolybdic acid solution (#27615, Nacalai Tesque) for 2 minutes at RT, the sections were stained in saturated picric acid containing 0.1% Sirius Red (#3306, Muto) for 90 minutes at RT. They were then rinsed in 0.5% hydrochloric acid solution for 1 minute, dehydrated in three changes of 100% ethanol and xylene, and finally coverslipped with Malinol (#2040, Muto).

Stained sections were examined microscopically in a non-blind manner by two pathologists. One was a veterinary pathologist with 3 years of experience, and the other was a certificated veterinary/toxicology pathologist with over 15 years of experience. The histopathological scores were established as follows: 0 - no change; 1 - minimal change; 2 - mild; 3 - moderate change; 4 - marked change; 5 - severe change. Sirius Red was also used to semi-quantify the extent of fibrosis.

#### Flow cytometry analysis

The liver sample was dissociated using gentleMACS according to the manufacturer’s instructions. Unless otherwise noted, PBS containing 2% fetal bovine serum was used as “Wash Buffer” thereafter. The washed samples were centrifuged (50×g, 4 °C for 3 min) to eliminate hepatocytes and were subjected to ACK Lysis Buffer. ACK buffer was prepared by adding 8,024 mg of NH_4_Cl (A2037, TCI, Japan), 10 mg of KHCO3 (166-03275, FUJIFILM Wako, Japan), and 3.772 mg of EDTA 2Na · 2H2O (6381-92-6, Dojindo Laboratories, Japan) into 1 L pure water. The samples were washed with the Wash Buffer three times and then the samples were subjected to flow cytometry analysis. Flow cytometric analysis was performed with FACSCelesta (BD Biosciences, USA), and data were analyzed with FlowJo software (Treestar). Proportions of NK cells (CD45+/CD3−/NK1.1+), B cells (CD45+/B220+), CD4+ T cells (CD45+/CD3+/B220−/CD4+), CD8+ T cells (CD45+/CD3+/B220−/CD4−/CD8a+), NKT cells (CD45+/CD3+/B220−/CD4−/CD8a−/NK1.1+), γδT cells (CD45+/CD3+/B220−/CD4−/CD8a−/gdTCR+), eosinophils (CD45+/CD11b+/Siglec-F+), neutrophils (CD45+/CD11b+/Siglec-F−/Ly6G+/Tim4−), monocytes (CD45+/CD11b+/Siglec-F−/Ly6G−/Tim4−/Ly6C+), monocyte-derived macrophages (CD45+/CD11b+/Siglec-F−/Ly6G−/Tim4−/Ly6C−), and Kupffer cells (CD45+/CD11b+/Siglec-F−/Ly6G−/Tim4+/Ly6C−) were acquired via flow cytometry. The antibodies used are listed in Supplementary Table S7.

#### Histopathological analysis with a deep learning model

The deep learning framework PyTorch version 2.0.1 was employed for this study. Deep learning and feature extraction tasks were carried out utilizing the FUJITSU Supercomputer PRIMEHPC FX1000 and FUJITSU Server PRIMERGY GX2570 (Wisteria/BDEC-01) at the Information Technology Center, The University of Tokyo. Python (version 3.11.3) with open-source packages was utilized for all analyses except for deep learning.

Pathological images of the liver were collected from the Open TG-GATEs (TGGATEs) database, along with corresponding pathological finding type annotations and individual rat information from TogoDB. These images were acquired at 20× magnification using Aperio ScanScope AT. The dataset preparation involved excluding compounds not sampled at all specified time points and limiting samples to the highest compound concentration. Subsequently, one control sample per condition and one image per individual were randomly selected, resulting in a dataset of 6508 whole slide images (WSI) for the liver. The construction of the finding-based model followed our previous work on expression learning based on pathological findings (3), with some differences such as varying image sizes and multi-label learning. The Open TG-GATEs dataset was split into Train1 and Test subsets in a 4:1 ratio, further split into Train2 and Validation subsets in a 4:1 ratio. Weak labels were generated through multi-label training using patches of pathological images and corresponding labels from the Train2 subset, focusing on eight pathological findings: proliferation bile duct, ground glass appearance, increased mitosis, inclusion body intracytoplasmic, deposit pigment, single cell necrosis, vacuolization cytoplasmic, and swelling. The model predicted the probability of these findings in images of the validation subset, repeated five times to apply resulting labels to Train1 subset patches. Patches were extracted based on predicted probabilities, and a new model was trained using these patches and corresponding probabilities for training of the finding-based model.

The formalin-fixed liver specimens underwent standard processing procedures, including slicing into 4 μm sections, staining with H&E, and visualization using ZEISS Axio Scan.Z1. Subsequently, the WSI were partitioned into patches of 256×256 pixels, and feature extraction was performed on each patch using the aforementioned model. After extracting features from the fourth layer, one class support vector machine (sklearn, version 1.4.1) was employed to classify whether each patch was anomalous or not. Additionally, dimensionality reduction of the features was achieved through PCA and visualized. Details are available on our GitHub repository (https://github.com/mizuno-group/MDAdrinkingwater).

## Supplementary Figures and Tables

**Fig. S1.**
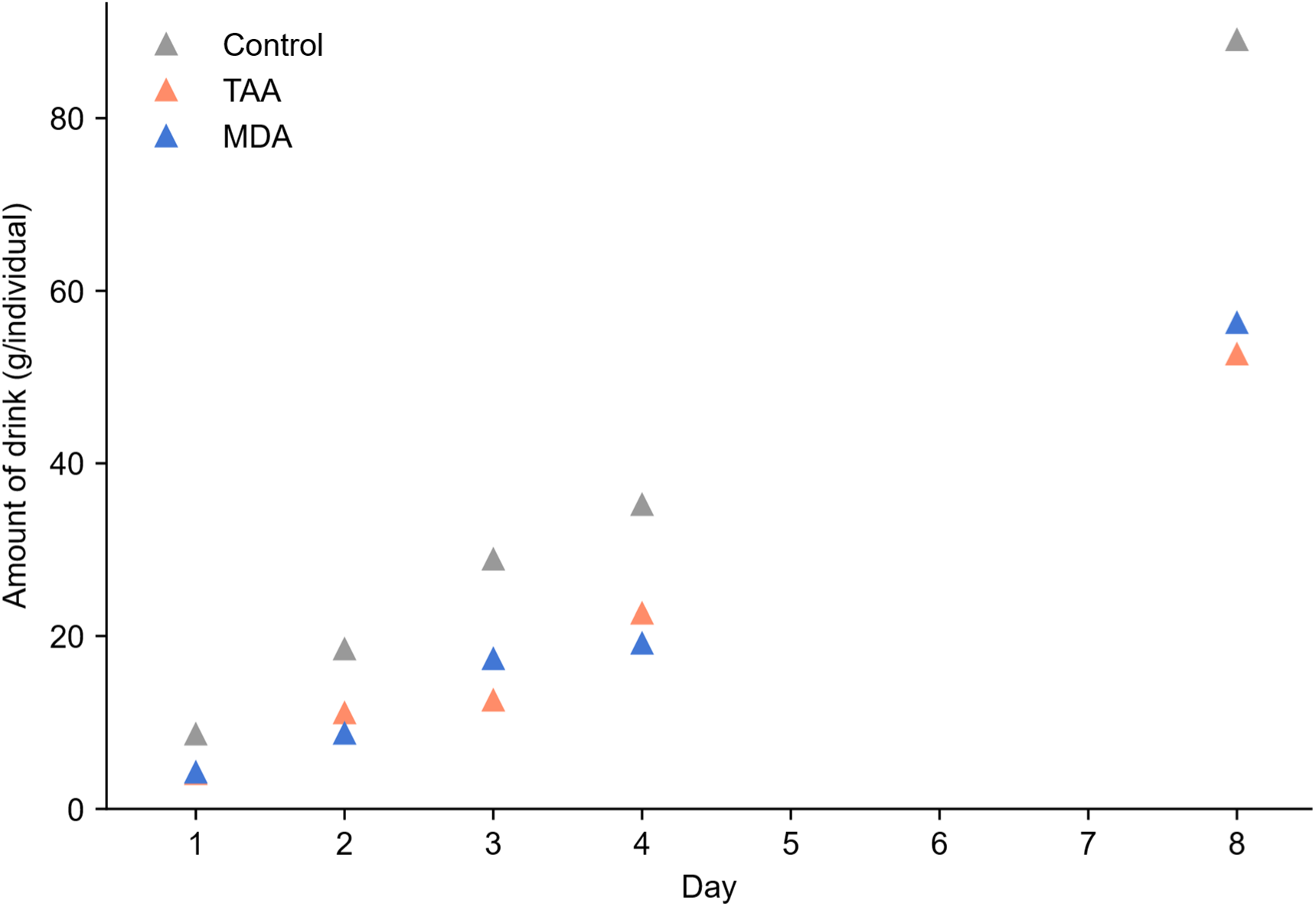
Cumulative estimates of water consumption in Cohort 2. Cumulative bottle weight loss from Day 0 to each sampling endpoint was normalized by the number of animals in the corresponding cage or cages. Separate cages were used for different endpoints. These estimates were not corrected for evaporation or leakage. Average daily intake over the initial intervals ending on Days 1–4 is summarized in Table 2.

**Fig. S2.**
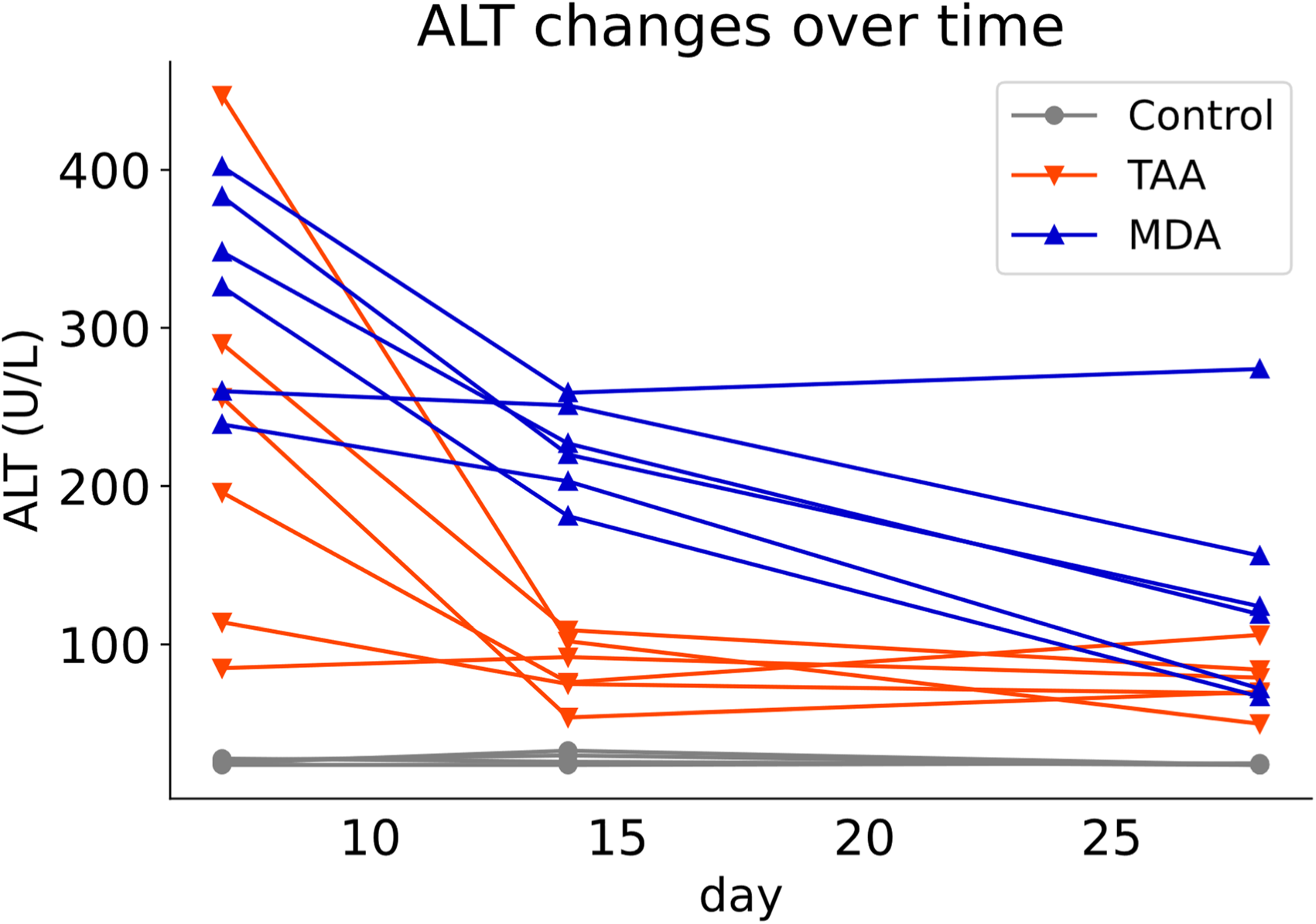
Plasma ALT changes during MDA-DW and TAA-DW administration in Cohort 1. Time-course changes in plasma alanine aminotransferase (ALT) levels are shown. Each line represents an individual animal followed through Day 28 (n = 4 controls, n = 6 TAA-DW, and n = 6 MDA-DW).

**Fig. S3.**
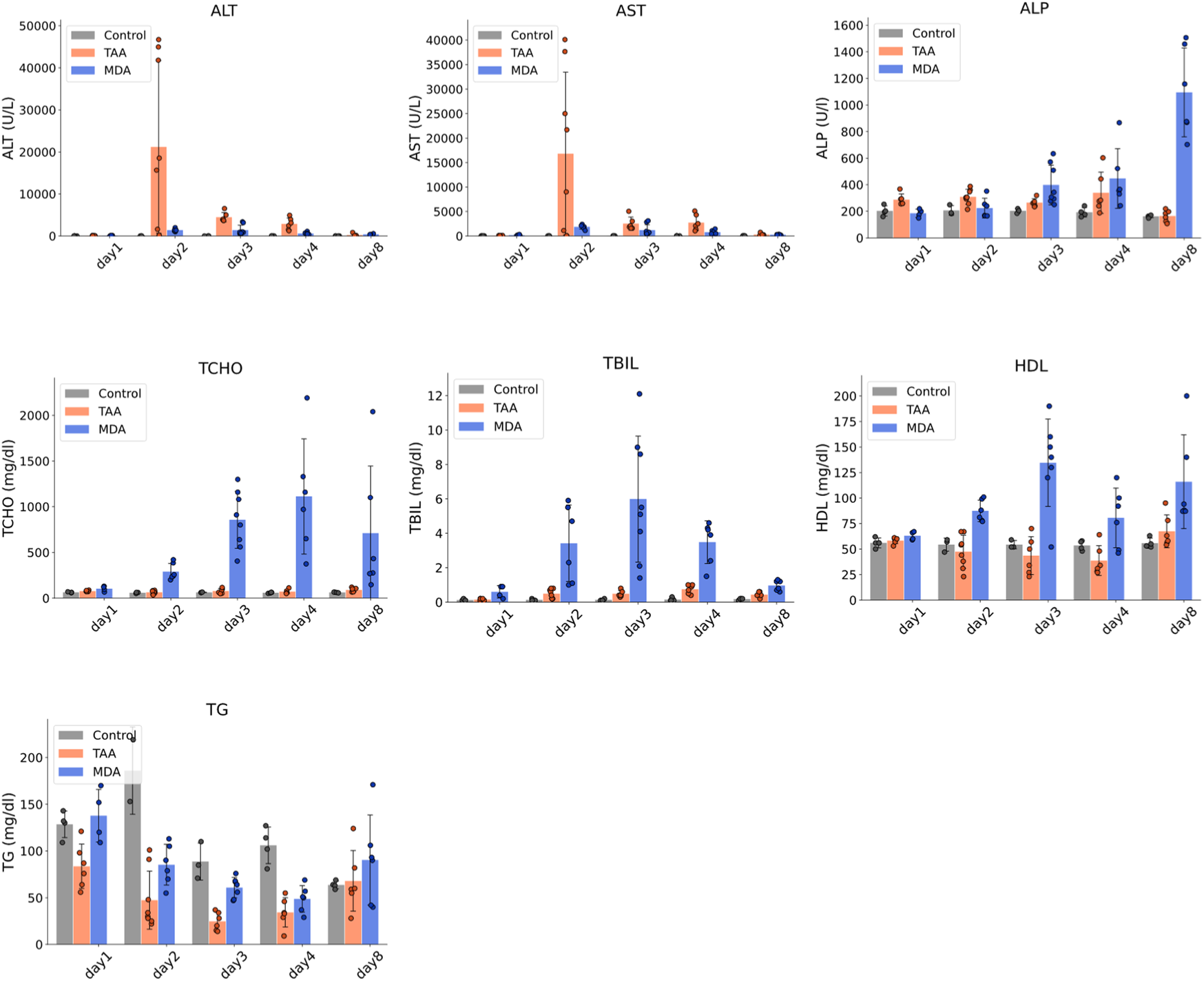
Plasma biochemical profiles in Cohort 2. Time-course changes in plasma biochemical values during the first 8 days of administration are shown. Bars represent mean ± SD, and dots represent individual animals. The x-axis is categorical and is not scaled according to elapsed time.

**Fig. S4.**
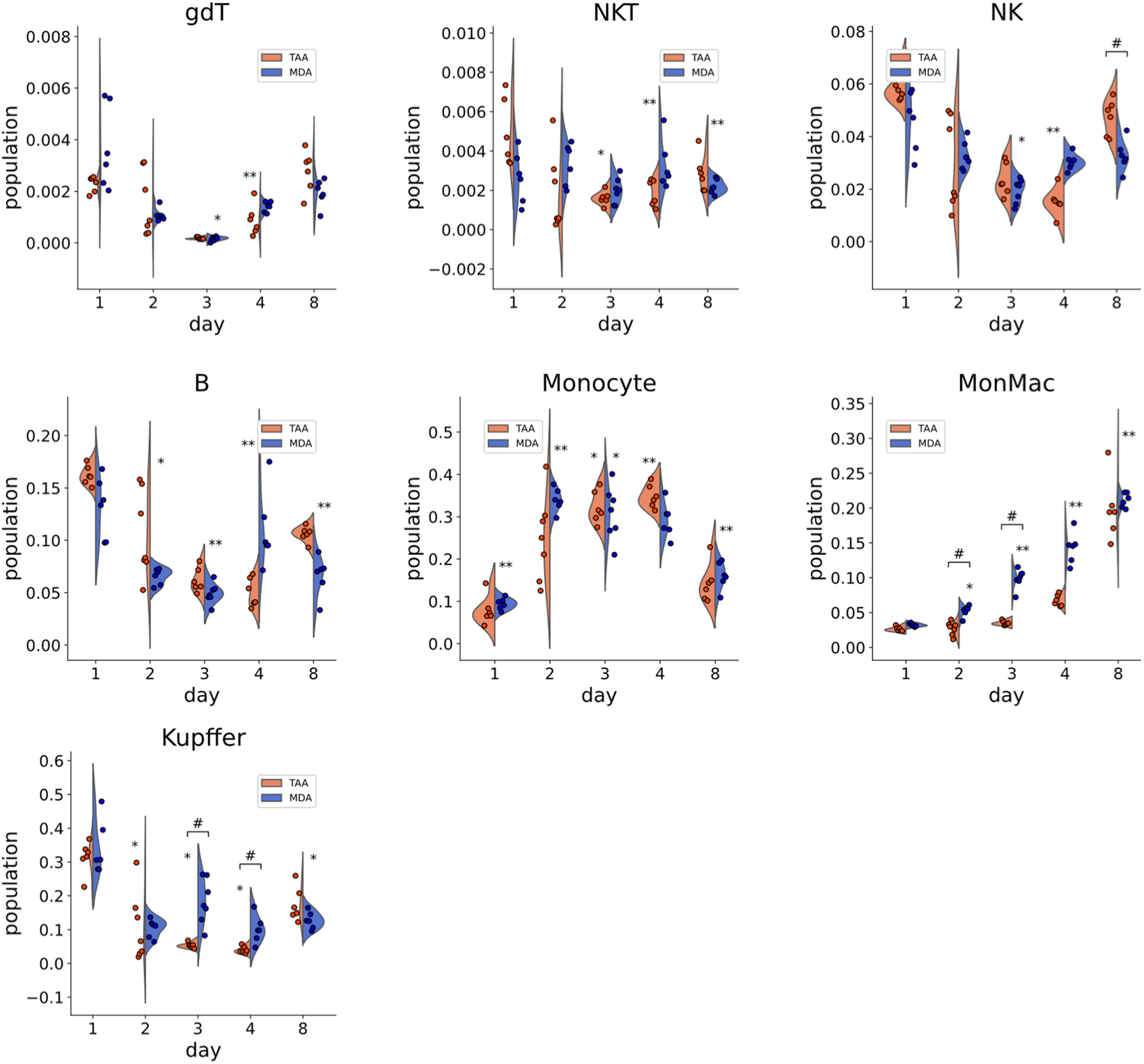
Additional hepatic immune-cell populations in Cohort 2. Violin plots show the proportions of immune-cell populations at each time point. Dots represent individual animals. The x-axis is categorical and is not scaled according to elapsed time. gdT, γδ T cells; MonMac, monocyte-derived macrophages; NKT, natural killer T cells; NK, natural killer cells. Statistical analysis used Kruskal–Wallis tests with Benjamini–Hochberg correction across all 55 population–time-point tests in Fig. 4B and Fig. S4, followed, where the adjusted P value was below 0.05, by Dunn’s pairwise tests with Holm correction within each population–time-point set. Asterisks indicate differences from the concurrent control group, which is included in the statistical analysis but not displayed. Hash marks indicate differences between MDA-DW and TAA-DW. One, two, and three symbols indicate Holm-adjusted P values < 0.05, < 0.01, and < 0.001, respectively.

**Fig. S5.**
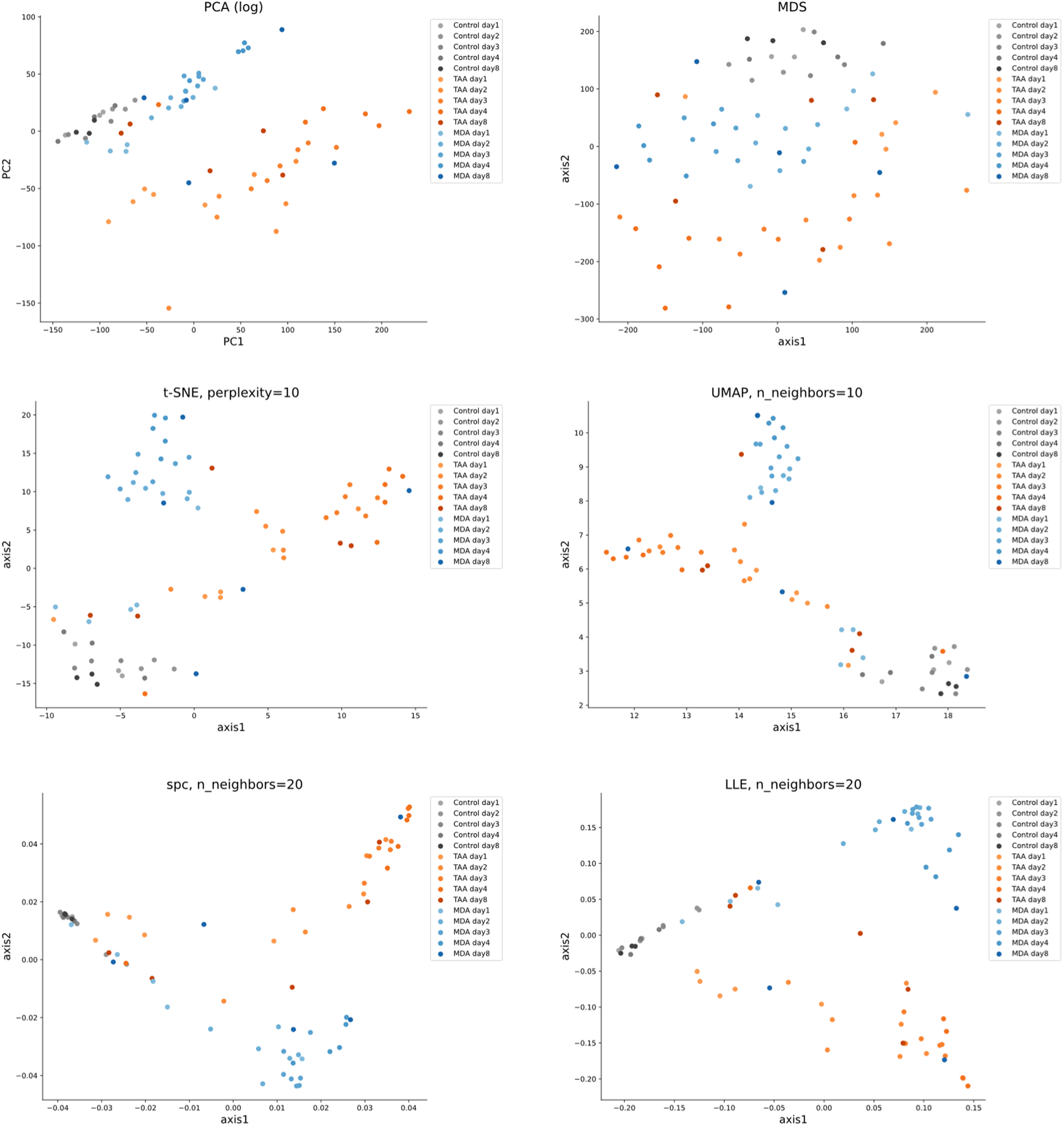
Individual dimensionality-reduction maps used for META visualization of transcriptome data. Transcriptome profiles were embedded in two dimensions using each dimensionality-reduction method included in META visualization. Values shown in the panel titles indicate the specified hyperparameter values. PCA, principal component analysis; MDS, multidimensional scaling; t-SNE, t-distributed stochastic neighbor embedding; UMAP, uniform manifold approximation and projection; spc, spectral embedding; LLE, locally linear embedding.

**Fig. S6.**
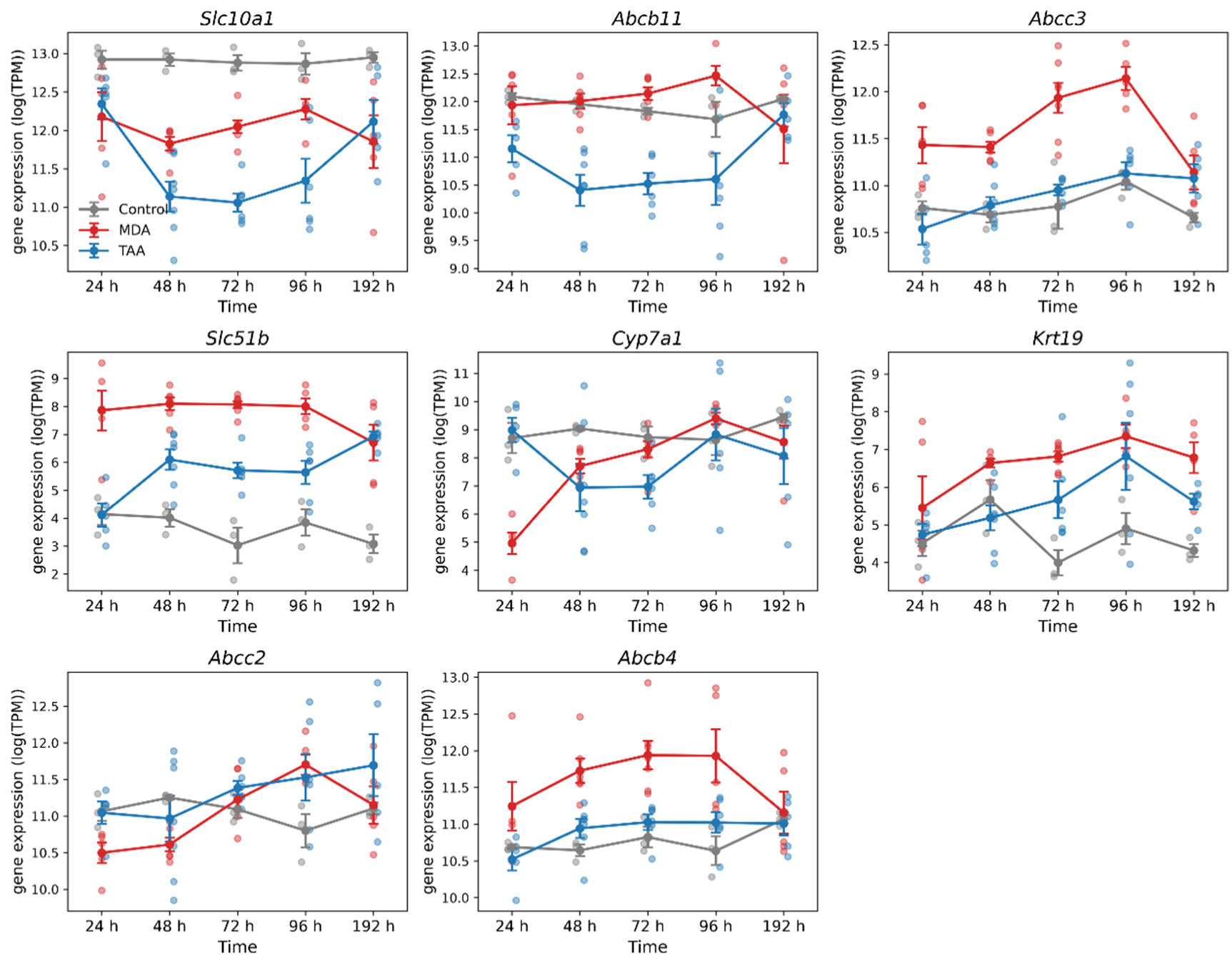
Expression profiles of selected genes related to bile acid homeostasis and biliary epithelial cells. Liver RNA-seq expression profiles of *Slc10a1*, *Abcb11*, *Abcc3*, *Slc51b*, *Cyp7a1*, *Krt19*, *Abcc2*, and *Abcb4* are shown for Cohort 2 on Days 1, 2, 3, 4, and 8. Dots represent individual animals; lines and error bars show the mean ± SEM of log₂-transformed scaled TPM. Sample sizes were 3 to 4 by group and time point. Lines connect group means from separate groups of animals sampled at each time point. The x-axis is categorical and is not scaled according to elapsed time. Values are presented descriptively.

---

**Supplementary Table S1:**
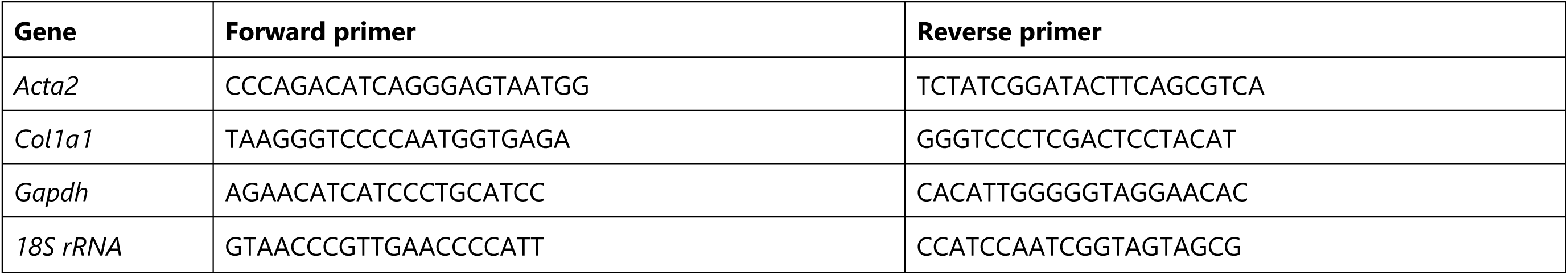
Sequences of qPCR primers.

**Supplementary Table S2:**
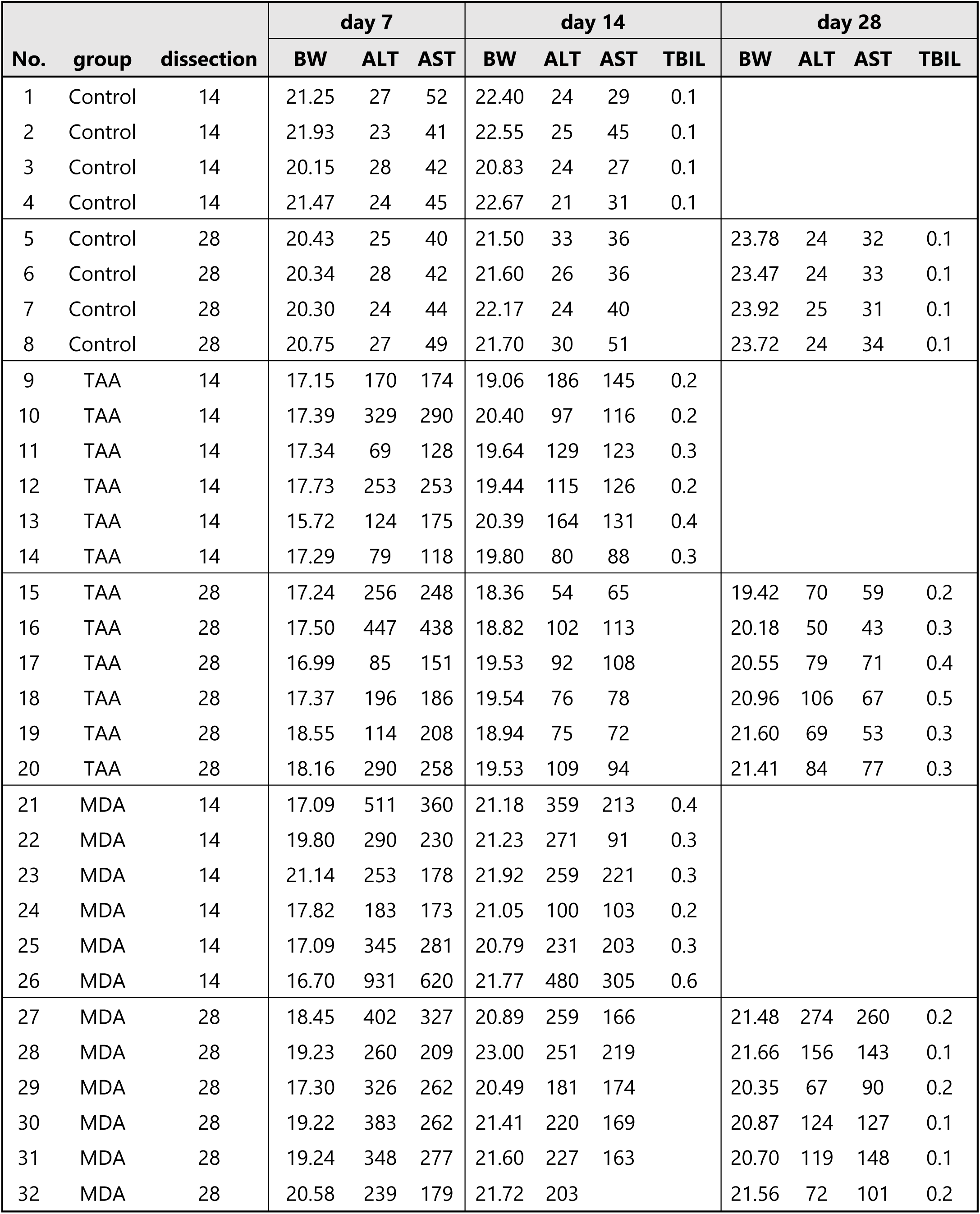
Blood biochemical values (Cohort 1)

| No. | group | dissection | day 7 |  |  | day 14 |  |  |  | day 28 |  |  |  |
| --- | --- | --- | --- | --- | --- | --- | --- | --- | --- | --- | --- | --- | --- |
|  |  |  | BW | ALT | AST | BW | ALT | AST | TBIL | BW | ALT | AST | TBIL |
| 1 | Control | 14 | 21.25 | 27 | 52 | 22.40 | 24 | 29 | 0.1 |  |  |  |  |
| 2 | Control | 14 | 21.93 | 23 | 41 | 22.55 | 25 | 45 | 0.1 |  |  |  |  |
| 3 | Control | 14 | 20.15 | 28 | 42 | 20.83 | 24 | 27 | 0.1 |  |  |  |  |
| 4 | Control | 14 | 21.47 | 24 | 45 | 22.67 | 21 | 31 | 0.1 |  |  |  |  |
| 5 | Control | 28 | 20.43 | 25 | 40 | 21.50 | 33 | 36 |  | 23.78 | 24 | 32 | 0.1 |
| 6 | Control | 28 | 20.34 | 28 | 42 | 21.60 | 26 | 36 |  | 23.47 | 24 | 33 | 0.1 |
| 7 | Control | 28 | 20.30 | 24 | 44 | 22.17 | 24 | 40 |  | 23.92 | 25 | 31 | 0.1 |
| 8 | Control | 28 | 20.75 | 27 | 49 | 21.70 | 30 | 51 |  | 23.72 | 24 | 34 | 0.1 |
| 9 | TAA | 14 | 17.15 | 170 | 174 | 19.06 | 186 | 145 | 0.2 |  |  |  |  |
| 10 | TAA | 14 | 17.39 | 329 | 290 | 20.40 | 97 | 116 | 0.2 |  |  |  |  |
| 11 | TAA | 14 | 17.34 | 69 | 128 | 19.64 | 129 | 123 | 0.3 |  |  |  |  |
| 12 | TAA | 14 | 17.73 | 253 | 253 | 19.44 | 115 | 126 | 0.2 |  |  |  |  |
| 13 | TAA | 14 | 15.72 | 124 | 175 | 20.39 | 164 | 131 | 0.4 |  |  |  |  |
| 14 | TAA | 14 | 17.29 | 79 | 118 | 19.80 | 80 | 88 | 0.3 |  |  |  |  |
| 15 | TAA | 28 | 17.24 | 256 | 248 | 18.36 | 54 | 65 |  | 19.42 | 70 | 59 | 0.2 |
| 16 | TAA | 28 | 17.50 | 447 | 438 | 18.82 | 102 | 113 |  | 20.18 | 50 | 43 | 0.3 |
| 17 | TAA | 28 | 16.99 | 85 | 151 | 19.53 | 92 | 108 |  | 20.55 | 79 | 71 | 0.4 |
| 18 | TAA | 28 | 17.37 | 196 | 186 | 19.54 | 76 | 78 |  | 20.96 | 106 | 67 | 0.5 |
| 19 | TAA | 28 | 18.55 | 114 | 208 | 18.94 | 75 | 72 |  | 21.60 | 69 | 53 | 0.3 |
| 20 | TAA | 28 | 18.16 | 290 | 258 | 19.53 | 109 | 94 |  | 21.41 | 84 | 77 | 0.3 |
| 21 | MDA | 14 | 17.09 | 511 | 360 | 21.18 | 359 | 213 | 0.4 |  |  |  |  |
| 22 | MDA | 14 | 19.80 | 290 | 230 | 21.23 | 271 | 91 | 0.3 |  |  |  |  |
| 23 | MDA | 14 | 21.14 | 253 | 178 | 21.92 | 259 | 221 | 0.3 |  |  |  |  |
| 24 | MDA | 14 | 17.82 | 183 | 173 | 21.05 | 100 | 103 | 0.2 |  |  |  |  |
| 25 | MDA | 14 | 17.09 | 345 | 281 | 20.79 | 231 | 203 | 0.3 |  |  |  |  |
| 26 | MDA | 14 | 16.70 | 931 | 620 | 21.77 | 480 | 305 | 0.6 |  |  |  |  |
| 27 | MDA | 28 | 18.45 | 402 | 327 | 20.89 | 259 | 166 |  | 21.48 | 274 | 260 | 0.2 |
| 28 | MDA | 28 | 19.23 | 260 | 209 | 23.00 | 251 | 219 |  | 21.66 | 156 | 143 | 0.1 |
| 29 | MDA | 28 | 17.30 | 326 | 262 | 20.49 | 181 | 174 |  | 20.35 | 67 | 90 | 0.2 |
| 30 | MDA | 28 | 19.22 | 383 | 262 | 21.41 | 220 | 169 |  | 20.87 | 124 | 127 | 0.1 |
| 31 | MDA | 28 | 19.24 | 348 | 277 | 21.60 | 227 | 163 |  | 20.70 | 119 | 148 | 0.1 |
| 32 | MDA | 28 | 20.58 | 239 | 179 | 21.72 | 203 |  |  | 21.56 | 72 | 101 | 0.2 |

**Supplementary Table S3.**
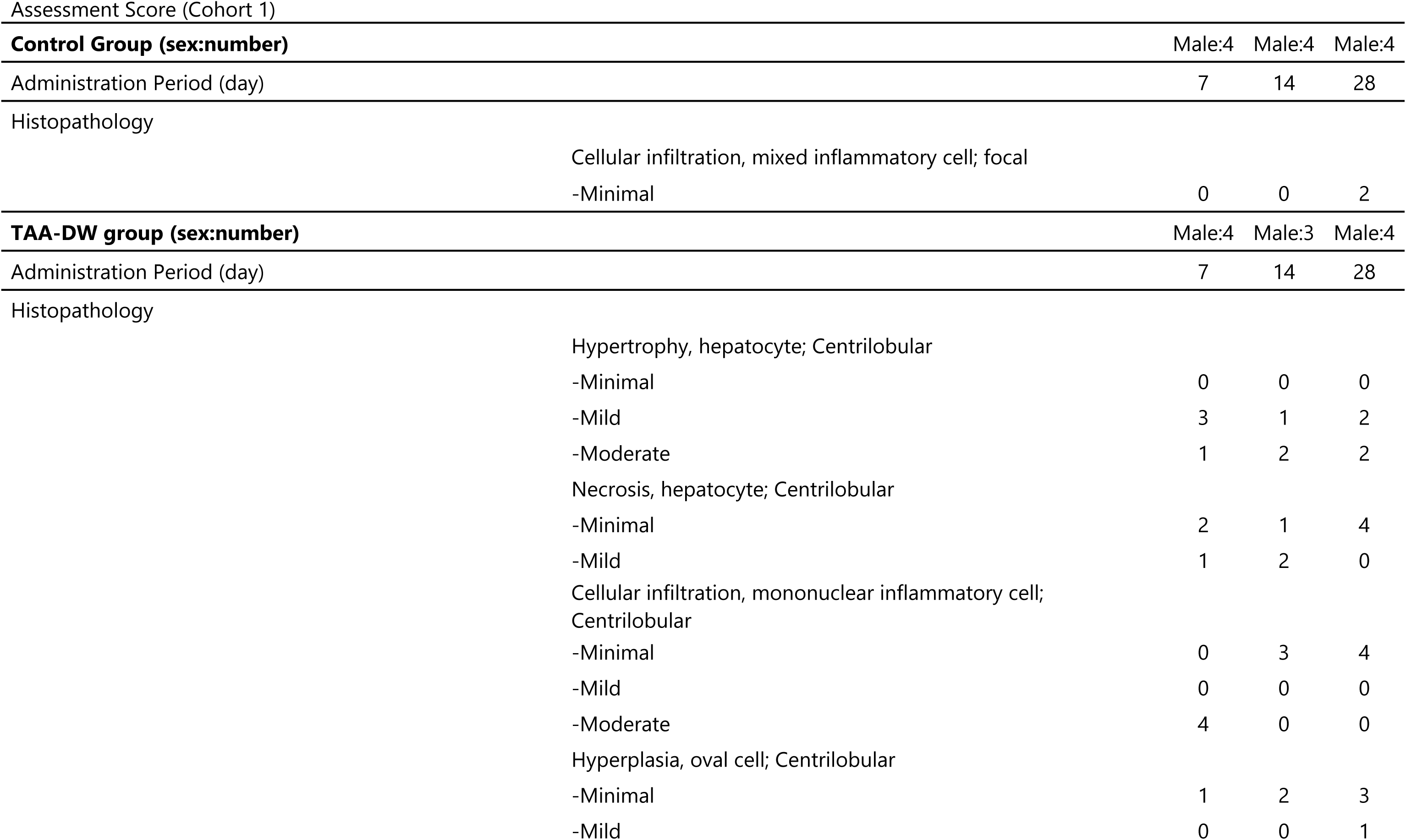
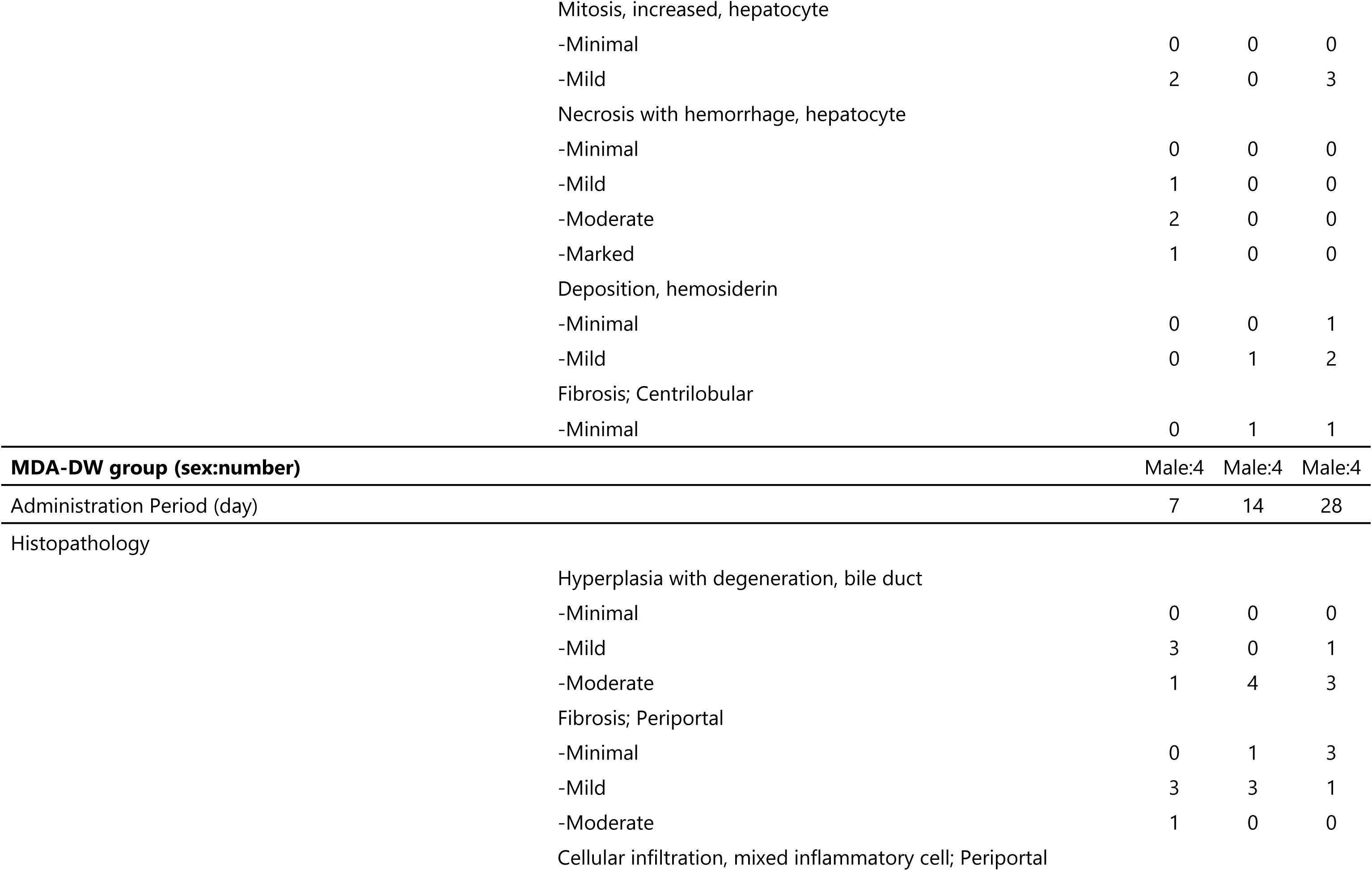
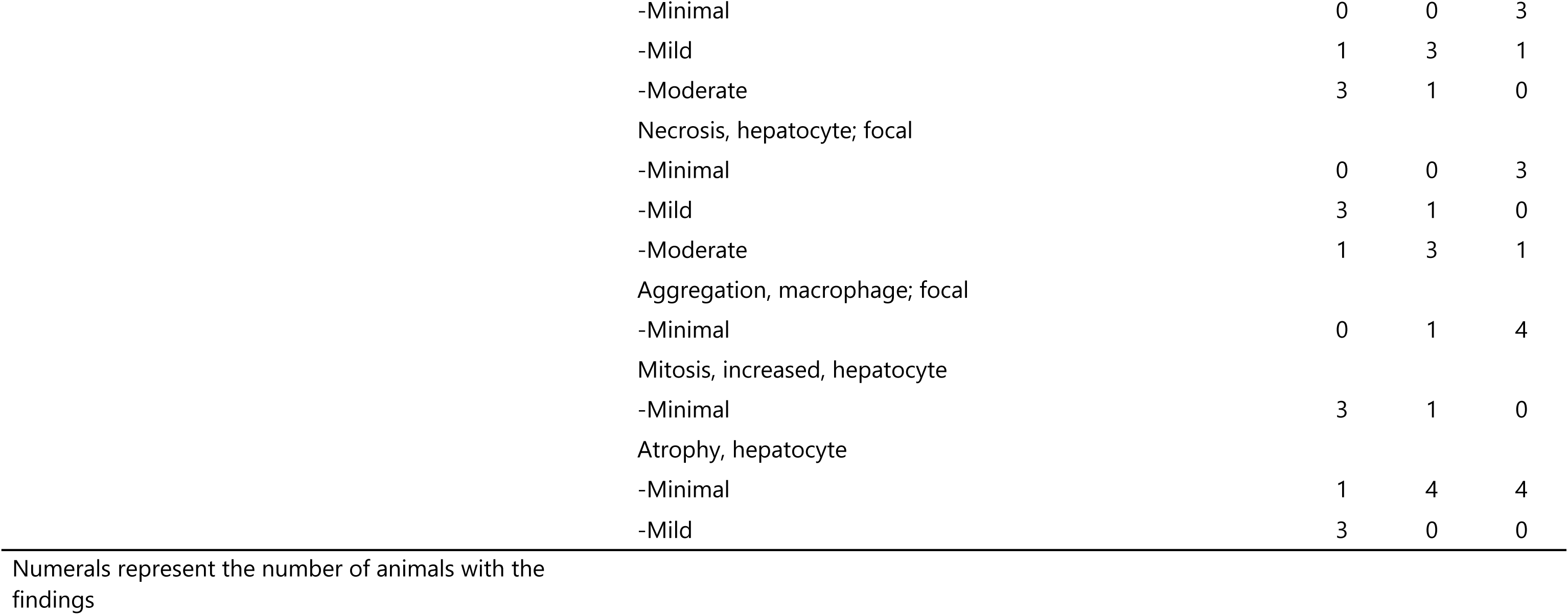
(a): Histopathological Liver.

**Supplementary Table S3.** (b): Histopathological scoring (control)

|  |  |  |  |  |  |  |  |  |  |  |  |  |
| --- | --- | --- | --- | --- | --- | --- | --- | --- | --- | --- | --- | --- |
| Control |  |  |  |  |  |  |  |  |  |  |  |  |
| Animal Number | 1 | 2 | 3 | 4 | 5 | 6 | 7 | 8 | 9 | 10 | 11 | 12 |
|  | N | N | N | *N | *N | N | N | N | *N | A | N | A |
| Cellular infiltration, mixed cell, focal | • | • | • | • | • | • | • | • | • | 1 | • | 1 |

\*: artifact
N=Normal; A=Abnormal; • =Not Recorded
1 : minimal

**Supplementary Table S3.**
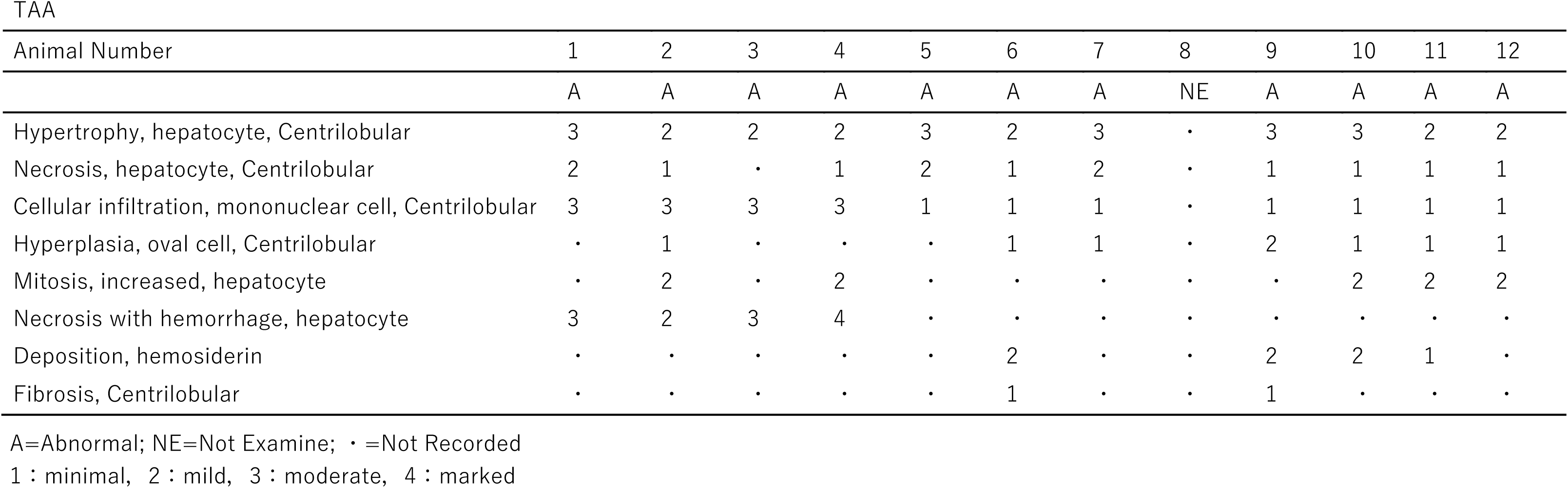
(c): Histopathological scoring (TAA)

| Animal Number | 1 | 2 | 3 | 4 | 5 | 6 | 7 | 8 | 9 | 10 | 11 | 12 |
| --- | --- | --- | --- | --- | --- | --- | --- | --- | --- | --- | --- | --- |
|  | A | A | A | A | A | A | A | NE | A | A | A | A |
| Hypertrophy, hepatocyte, Centrilobular | 3 | 2 | 2 | 2 | 3 | 2 | 3 | • | 3 | 3 | 2 | 2 |
| Necrosis, hepatocyte, Centrilobular | 2 | 1 | • | 1 | 2 | 1 | 2 | • | 1 | 1 | 1 | 1 |
| Cellular infiltration, mononuclear cell, Centrilobular | 3 | 3 | 3 | 3 | 1 | 1 | 1 | • | 1 | 1 | 1 | 1 |
| Hyperplasia, oval cell, Centrilobular | • | 1 | • | • | • | 1 | 1 | • | 2 | 1 | 1 | 1 |
| Mitosis, increased, hepatocyte | • | 2 | • | 2 | • | • | • | • | • | 2 | 2 | 2 |
| Necrosis with hemorrhage, hepatocyte | 3 | 2 | 3 | 4 | • | • | • | • | • | • | • | • |
| Deposition, hemosiderin | • | • | • | • | • | 2 | • | • | 2 | 2 | 1 | • |
| Fibrosis, Centrilobular | • | • | • | • | • | 1 | • | • | 1 | • | • | • |

A=Abnormal; NE=Not Examine; • =Not Recorded
1 : minimal, 2 : mild, 3 : moderate, 4 : marked

**Supplementary Table S3.** (d): Histopathological scoring (MDA)

| Animal Number | 1 | 2 | 3 | 4 | 5 | 6 | 7 | 8 | 9 | 10 | 11 | 12 |
| --- | --- | --- | --- | --- | --- | --- | --- | --- | --- | --- | --- | --- |
|  | A | A | A | A | A | A | A | A | A | *A | A | A |
| Hyperplasia with degeneration, bile duct | 2 | 3 | 2 | 2 | 3 | 3 | 3 | 3 | 2 | 3 | 3 | 3 |
| Fibrosis, Periportal | 2 | 2 | 2 | 3 | 2 | 1 | 2 | 2 | 1 | 1 | 2 | 1 |
| Cellular infiltration, mixed cell, Periportal | 3 | 3 | 3 | 2 | 2 | 2 | 2 | 3 | 2 | 2 | 1 | 1 |
| Necrosis, hepatocyte, focal | 2 | 2 | 3 | 2 | 3 | 3 | 2 | 3 | 1 | 3 | 1 | 1 |
| Aggregation, macrophage, focal | • | • | • | • | • | • | 2 | • | 2 | 2 | 2 | 2 |
| Mitosis, increased, hepatocyte | 1 | 1 | • | 1 | • | 1 | • | • | • | • | • | • |
| Atrophy, hepatocyte | 2 | 2 | 2 | 1 | 1 | 1 | 1 | 1 | 1 | 1 | 1 | 1 |

\*: artifact
A=Abnormal; • =Not Recorded
1 : minimal, 2 : mild, 3 : moderate

**Supplementary Table S4a:**
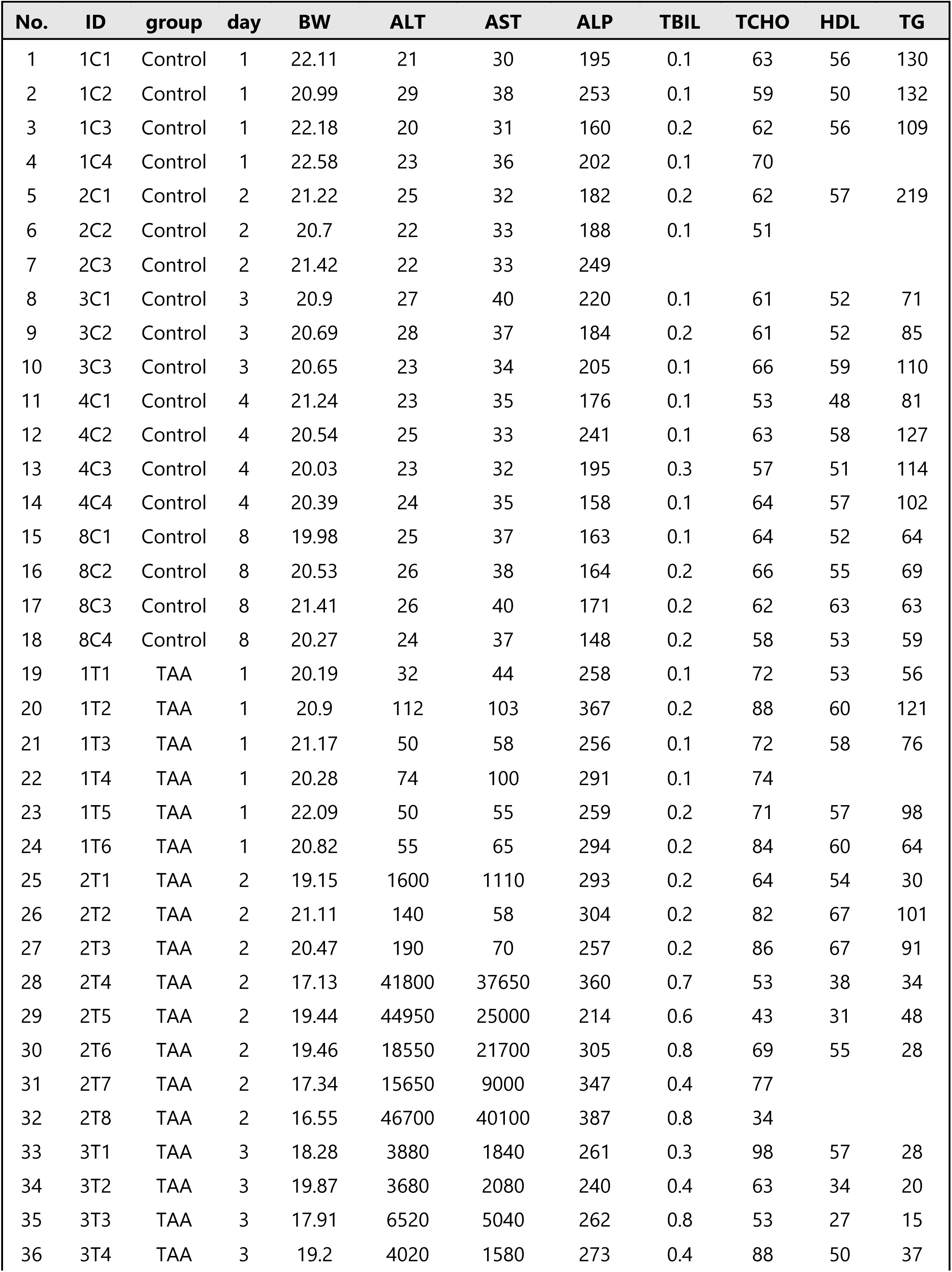
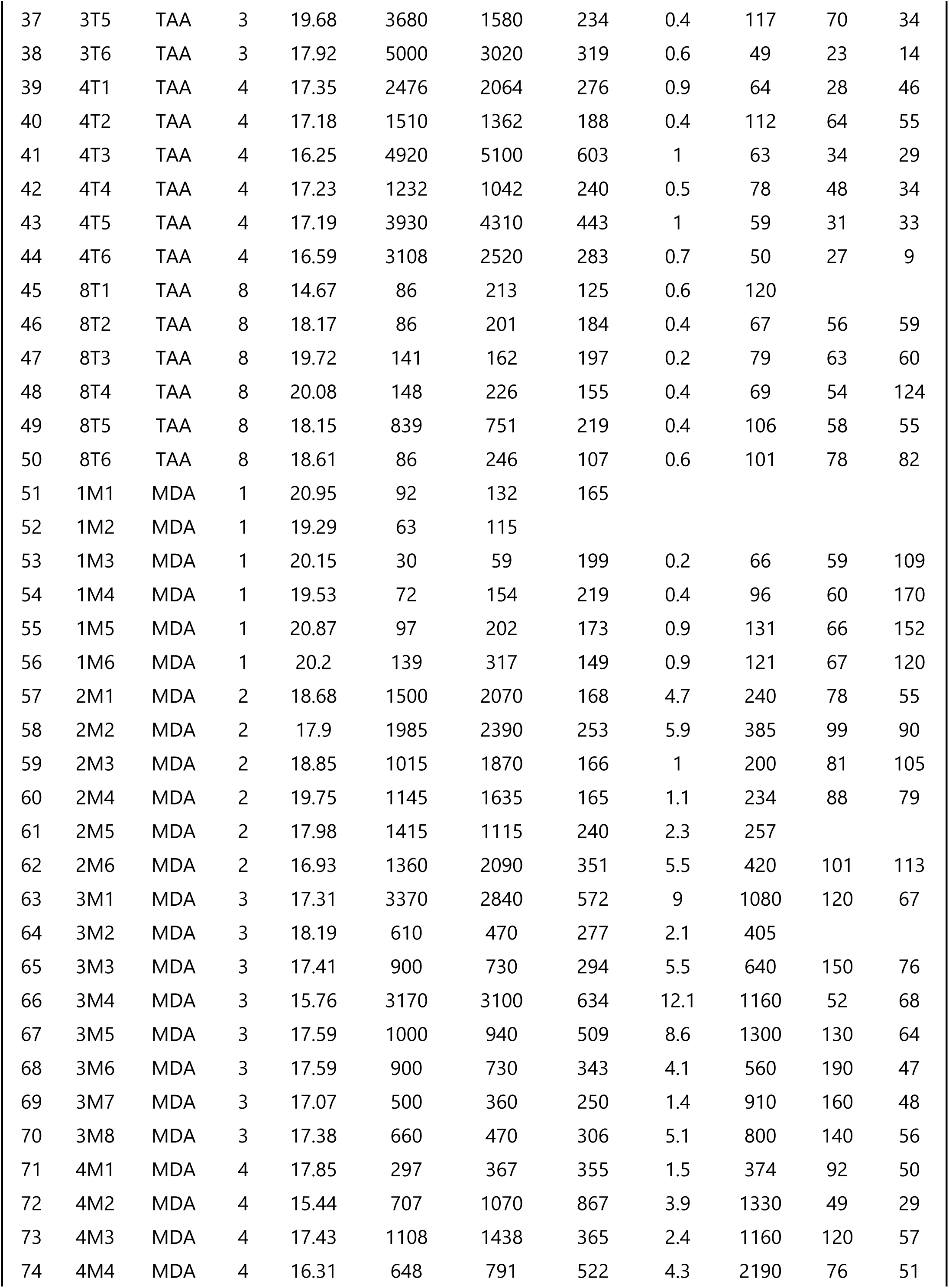
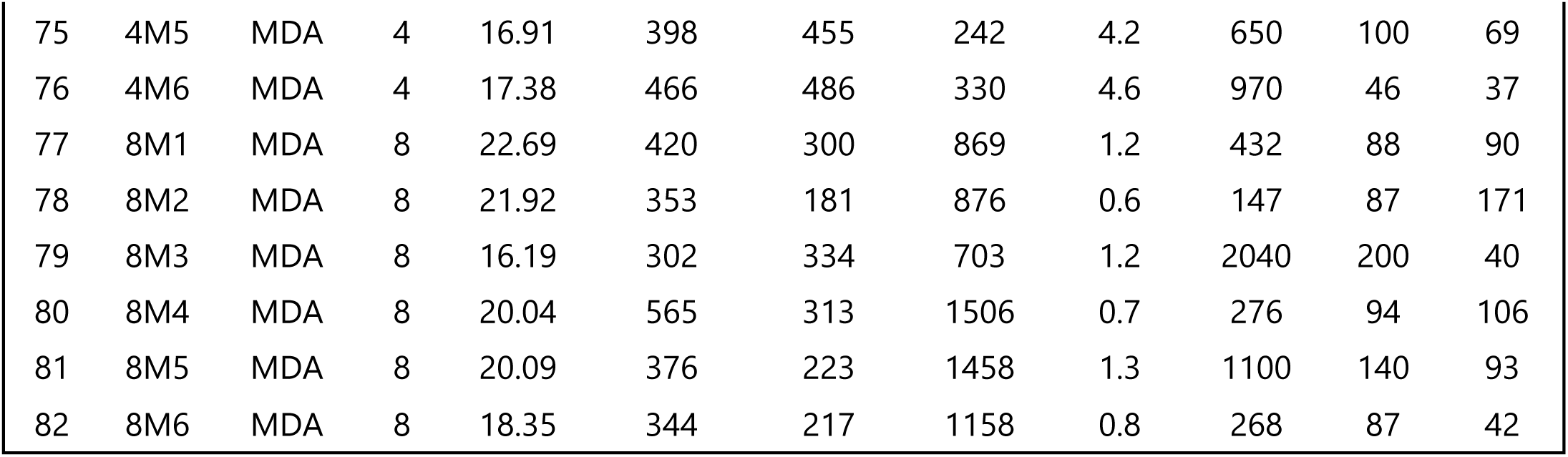
Blood biochemical values (Cohort 2)

| No. | ID | group | day | BW | ALT | AST | ALP | TBIL | TCHO | HDL | TG |
| --- | --- | --- | --- | --- | --- | --- | --- | --- | --- | --- | --- |
| 1 | 1C1 | Control | 1 | 22.11 | 21 | 30 | 195 | 0.1 | 63 | 56 | 130 |
| 2 | 1C2 | Control | 1 | 20.99 | 29 | 38 | 253 | 0.1 | 59 | 50 | 132 |
| 3 | 1C3 | Control | 1 | 22.18 | 20 | 31 | 160 | 0.2 | 62 | 56 | 109 |
| 4 | 1C4 | Control | 1 | 22.58 | 23 | 36 | 202 | 0.1 | 70 |  |  |
| 5 | 2C1 | Control | 2 | 21.22 | 25 | 32 | 182 | 0.2 | 62 | 57 | 219 |
| 6 | 2C2 | Control | 2 | 20.7 | 22 | 33 | 188 | 0.1 | 51 |  |  |
| 7 | 2C3 | Control | 2 | 21.42 | 22 | 33 | 249 |  |  |  |  |
| 8 | 3C1 | Control | 3 | 20.9 | 27 | 40 | 220 | 0.1 | 61 | 52 | 71 |
| 9 | 3C2 | Control | 3 | 20.69 | 28 | 37 | 184 | 0.2 | 61 | 52 | 85 |
| 10 | 3C3 | Control | 3 | 20.65 | 23 | 34 | 205 | 0.1 | 66 | 59 | 110 |
| 11 | 4C1 | Control | 4 | 21.24 | 23 | 35 | 176 | 0.1 | 53 | 48 | 81 |
| 12 | 4C2 | Control | 4 | 20.54 | 25 | 33 | 241 | 0.1 | 63 | 58 | 127 |
| 13 | 4C3 | Control | 4 | 20.03 | 23 | 32 | 195 | 0.3 | 57 | 51 | 114 |
| 14 | 4C4 | Control | 4 | 20.39 | 24 | 35 | 158 | 0.1 | 64 | 57 | 102 |
| 15 | 8C1 | Control | 8 | 19.98 | 25 | 37 | 163 | 0.1 | 64 | 52 | 64 |
| 16 | 8C2 | Control | 8 | 20.53 | 26 | 38 | 164 | 0.2 | 66 | 55 | 69 |
| 17 | 8C3 | Control | 8 | 21.41 | 26 | 40 | 171 | 0.2 | 62 | 63 | 63 |
| 18 | 8C4 | Control | 8 | 20.27 | 24 | 37 | 148 | 0.2 | 58 | 53 | 59 |
| 19 | 1T1 | TAA | 1 | 20.19 | 32 | 44 | 258 | 0.1 | 72 | 53 | 56 |
| 20 | 1T2 | TAA | 1 | 20.9 | 112 | 103 | 367 | 0.2 | 88 | 60 | 121 |
| 21 | 1T3 | TAA | 1 | 21.17 | 50 | 58 | 256 | 0.1 | 72 | 58 | 76 |
| 22 | 1T4 | TAA | 1 | 20.28 | 74 | 100 | 291 | 0.1 | 74 |  |  |
| 23 | 1T5 | TAA | 1 | 22.09 | 50 | 55 | 259 | 0.2 | 71 | 57 | 98 |
| 24 | 1T6 | TAA | 1 | 20.82 | 55 | 65 | 294 | 0.2 | 84 | 60 | 64 |
| 25 | 2T1 | TAA | 2 | 19.15 | 1600 | 1110 | 293 | 0.2 | 64 | 54 | 30 |
| 26 | 2T2 | TAA | 2 | 21.11 | 140 | 58 | 304 | 0.2 | 82 | 67 | 101 |
| 27 | 2T3 | TAA | 2 | 20.47 | 190 | 70 | 257 | 0.2 | 86 | 67 | 91 |
| 28 | 2T4 | TAA | 2 | 17.13 | 41800 | 37650 | 360 | 0.7 | 53 | 38 | 34 |
| 29 | 2T5 | TAA | 2 | 19.44 | 44950 | 25000 | 214 | 0.6 | 43 | 31 | 48 |
| 30 | 2T6 | TAA | 2 | 19.46 | 18550 | 21700 | 305 | 0.8 | 69 | 55 | 28 |
| 31 | 2T7 | TAA | 2 | 17.34 | 15650 | 9000 | 347 | 0.4 | 77 |  |  |
| 32 | 2T8 | TAA | 2 | 16.55 | 46700 | 40100 | 387 | 0.8 | 34 |  |  |
| 33 | 3T1 | TAA | 3 | 18.28 | 3880 | 1840 | 261 | 0.3 | 98 | 57 | 28 |
| 34 | 3T2 | TAA | 3 | 19.87 | 3680 | 2080 | 240 | 0.4 | 63 | 34 | 20 |
| 35 | 3T3 | TAA | 3 | 17.91 | 6520 | 5040 | 262 | 0.8 | 53 | 27 | 15 |
| 36 | 3T4 | TAA | 3 | 19.2 | 4020 | 1580 | 273 | 0.4 | 88 | 50 | 37 |
| 37 | 3T5 | TAA | 3 | 19.68 | 3680 | 1580 | 234 | 0.4 | 117 | 70 | 34 |
| 38 | 3T6 | TAA | 3 | 17.92 | 5000 | 3020 | 319 | 0.6 | 49 | 23 | 14 |
| 39 | 4T1 | TAA | 4 | 17.35 | 2476 | 2064 | 276 | 0.9 | 64 | 28 | 46 |
| 40 | 4T2 | TAA | 4 | 17.18 | 1510 | 1362 | 188 | 0.4 | 112 | 64 | 55 |
| 41 | 4T3 | TAA | 4 | 16.25 | 4920 | 5100 | 603 | 1 | 63 | 34 | 29 |
| 42 | 4T4 | TAA | 4 | 17.23 | 1232 | 1042 | 240 | 0.5 | 78 | 48 | 34 |
| 43 | 4T5 | TAA | 4 | 17.19 | 3930 | 4310 | 443 | 1 | 59 | 31 | 33 |
| 44 | 4T6 | TAA | 4 | 16.59 | 3108 | 2520 | 283 | 0.7 | 50 | 27 | 9 |
| 45 | 8T1 | TAA | 8 | 14.67 | 86 | 213 | 125 | 0.6 | 120 |  |  |
| 46 | 8T2 | TAA | 8 | 18.17 | 86 | 201 | 184 | 0.4 | 67 | 56 | 59 |
| 47 | 8T3 | TAA | 8 | 19.72 | 141 | 162 | 197 | 0.2 | 79 | 63 | 60 |
| 48 | 8T4 | TAA | 8 | 20.08 | 148 | 226 | 155 | 0.4 | 69 | 54 | 124 |
| 49 | 8T5 | TAA | 8 | 18.15 | 839 | 751 | 219 | 0.4 | 106 | 58 | 55 |
| 50 | 8T6 | TAA | 8 | 18.61 | 86 | 246 | 107 | 0.6 | 101 | 78 | 82 |
| 51 | 1M1 | MDA | 1 | 20.95 | 92 | 132 | 165 |  |  |  |  |
| 52 | 1M2 | MDA | 1 | 19.29 | 63 | 115 |  |  |  |  |  |
| 53 | 1M3 | MDA | 1 | 20.15 | 30 | 59 | 199 | 0.2 | 66 | 59 | 109 |
| 54 | 1M4 | MDA | 1 | 19.53 | 72 | 154 | 219 | 0.4 | 96 | 60 | 170 |
| 55 | 1M5 | MDA | 1 | 20.87 | 97 | 202 | 173 | 0.9 | 131 | 66 | 152 |
| 56 | 1M6 | MDA | 1 | 20.2 | 139 | 317 | 149 | 0.9 | 121 | 67 | 120 |
| 57 | 2M1 | MDA | 2 | 18.68 | 1500 | 2070 | 168 | 4.7 | 240 | 78 | 55 |
| 58 | 2M2 | MDA | 2 | 17.9 | 1985 | 2390 | 253 | 5.9 | 385 | 99 | 90 |
| 59 | 2M3 | MDA | 2 | 18.85 | 1015 | 1870 | 166 | 1 | 200 | 81 | 105 |
| 60 | 2M4 | MDA | 2 | 19.75 | 1145 | 1635 | 165 | 1.1 | 234 | 88 | 79 |
| 61 | 2M5 | MDA | 2 | 17.98 | 1415 | 1115 | 240 | 2.3 | 257 |  |  |
| 62 | 2M6 | MDA | 2 | 16.93 | 1360 | 2090 | 351 | 5.5 | 420 | 101 | 113 |
| 63 | 3M1 | MDA | 3 | 17.31 | 3370 | 2840 | 572 | 9 | 1080 | 120 | 67 |
| 64 | 3M2 | MDA | 3 | 18.19 | 610 | 470 | 277 | 2.1 | 405 |  |  |
| 65 | 3M3 | MDA | 3 | 17.41 | 900 | 730 | 294 | 5.5 | 640 | 150 | 76 |
| 66 | 3M4 | MDA | 3 | 15.76 | 3170 | 3100 | 634 | 12.1 | 1160 | 52 | 68 |
| 67 | 3M5 | MDA | 3 | 17.59 | 1000 | 940 | 509 | 8.6 | 1300 | 130 | 64 |
| 68 | 3M6 | MDA | 3 | 17.59 | 900 | 730 | 343 | 4.1 | 560 | 190 | 47 |
| 69 | 3M7 | MDA | 3 | 17.07 | 500 | 360 | 250 | 1.4 | 910 | 160 | 48 |
| 70 | 3M8 | MDA | 3 | 17.38 | 660 | 470 | 306 | 5.1 | 800 | 140 | 56 |
| 71 | 4M1 | MDA | 4 | 17.85 | 297 | 367 | 355 | 1.5 | 374 | 92 | 50 |
| 72 | 4M2 | MDA | 4 | 15.44 | 707 | 1070 | 867 | 3.9 | 1330 | 49 | 29 |
| 73 | 4M3 | MDA | 4 | 17.43 | 1108 | 1438 | 365 | 2.4 | 1160 | 120 | 57 |
| 74 | 4M4 | MDA | 4 | 16.31 | 648 | 791 | 522 | 4.3 | 2190 | 76 | 51 |
| 75 | 4M5 | MDA | 4 | 16.91 | 398 | 455 | 242 | 4.2 | 650 | 100 | 69 |
| 76 | 4M6 | MDA | 4 | 17.38 | 466 | 486 | 330 | 4.6 | 970 | 46 | 37 |
| 77 | 8M1 | MDA | 8 | 22.69 | 420 | 300 | 869 | 1.2 | 432 | 88 | 90 |
| 78 | 8M2 | MDA | 8 | 21.92 | 353 | 181 | 876 | 0.6 | 147 | 87 | 171 |
| 79 | 8M3 | MDA | 8 | 16.19 | 302 | 334 | 703 | 1.2 | 2040 | 200 | 40 |
| 80 | 8M4 | MDA | 8 | 20.04 | 565 | 313 | 1506 | 0.7 | 276 | 94 | 106 |
| 81 | 8M5 | MDA | 8 | 20.09 | 376 | 223 | 1458 | 1.3 | 1100 | 140 | 93 |
| 82 | 8M6 | MDA | 8 | 18.35 | 344 | 217 | 1158 | 0.8 | 268 | 87 | 42 |

**Supplementary Table S4b:** Blood biochemical values (Cohort 2)

| day | Control |  |  | TAA |  |  | MDA |  |  |
| --- | --- | --- | --- | --- | --- | --- | --- | --- | --- |
|  | BW | ALT | AST | BW | ALT | AST | BW | ALT | AST |
| 1 | 22.0±0.7 | 23.3±4.0 | 33.8±3.9 | 20.9±0.7 | 62.2±27.9 | 70.8±24.7 | 20.2±0.7 | 82.2±36.7 | 163.2±88.8 |
| 2 | 21.1±0.4 | 23.0±1.7 | 32.7±0.6 | 18.8±1.7 | 21197.5±20513.4 | 16836.0±16630.7 | 18.3±1.0 | 1403.3±336.8 | 1861.7±443.5 |
| 3 | 20.7±0.1 | 26.0±2.6 | 37.0±3.0 | 18.8±0.9 | 4463.3±1120.9 | 2523.3±1343.4 | 17.3±0.7 | 1388.8±1174.4 | 1205.0±1107.1 |
| 4 | 20.6±0.5 | 23.8±1.0 | 33.8±1.5 | 17.0±0.4 | 2862.7±1419.2 | 2733.0±1632.3 | 16.9±0.9 | 604.0±290.6 | 767.8±419.3 |
| 8 | 20.5±0.6 | 25.3±1.0 | 38.0±1.4 | 18.2±1.9 | 231.0±299.2 | 299.8±222.8 | 19.9±2.4 | 393.3±92.6 | 261.3±62.2 |

**Supplementary Table S5:** Result of rankproduct (MDA group)

| **GO** | **log rank product statistic** | **p value** | **adjusted p value** |
| --- | --- | --- | --- |
| GOBP_RESPONSE_TO_INTERLEUKIN_17 | 23.59441201 | 1.72E-08 | 8.93357E-05 |
| GOBP_ACUTE_PHASE_RESPONSE | 21.39718743 | 1.28E-07 | 0.000333502 |
| GOBP_POSITIVE_REGULATION_OF_LEUKOCYTE_ADHESION_TO_VASCULAR_ENDOTHELIAL_CELL | 20.50336956 | 2.89E-07 | 0.000501057 |
| GOBP_POSITIVE_REGULATION_OF_MONOCYTE_CHEMOTAXIS | 19.00944453 | 1.11E-06 | 0.001143545 |
| GOBP_TOLL_LIKE_RECEPTOR_2_SIGNALING_PATHWAY | 18.8916615 | 1.24E-06 | 0.001143545 |
| GOBP_INTERLEUKIN_1_MEDIATED_SIGNALING_PATHWAY | 18.82266862 | 1.32E-06 | 0.001143545 |
| GOBP_PROTEIN_N_LINKED_GLYCOSYLATION_VIA ASPARAGINE | 18.58377672 | 1.63E-06 | 0.001183214 |
| GOBP_NEGATIVE_REGULATION_OF_EXTRINSIC_APOPTOTIC_SIGNALING_PATHWAY_VIA_DEATH_DOMAIN_RECEPTORS | 18.4645133 | 1.82E-06 | 0.001183214 |
| GOBP_DENDRITIC_CELL_DIFFERENTIATION | 17.92885075 | 2.94E-06 | 0.001699703 |
| GOBP_T_HELPER_1_CELL_DIFFERENTIATION | 17.26202088 | 5.33E-06 | 0.002522101 |
| GOBP_FIBRINOLYSIS | 16.72306141 | 8.60E-06 | 0.003733369 |
| GOCC_COPI_COATED_VESICLE_MEMBRANE | 16.58094628 | 9.76E-06 | 0.003909202 |
| GOMF_MISFOLDED_PROTEIN_BINDING | 16.47659842 | 1.07E-05 | 0.003981687 |
| GOBP_POSITIVE_REGULATION_OF_PHAGOCYTOSIS_ENGULFMENT | 16.32898229 | 1.22E-05 | 0.004235156 |
| GOBP_REGULATION_OF_HETEROTYPIC_CELL_CELL_ADHESION | 15.42819574 | 2.70E-05 | 0.008270168 |
| GOBP_LEUKOCYTE_MIGRATION_INVOLVED_IN_INFLAMMATORY_RESPONSE | 15.28572009 | 3.06E-05 | 0.008684274 |
| GOBP_I_KAPPAB_PHOSPHORYLATION | 15.24587418 | 3.17E-05 | 0.008684274 |
| GOBP_ENDOPLASMIC_RETICULUM_TO_CYTOSOL_TRANSPORT | 14.7995492 | 4.68E-05 | 0.011614524 |
| GOBP_POSITIVE_REGULATION_OF_MACROPHAGE_CYTOKINE_PRODUCTION | 14.71111843 | 5.06E-05 | 0.011976983 |
| GOBP_ANTIFUNGAL_INNATE_IMMUNE_RESPONSE | 14.63301388 | 5.42E-05 | 0.012264544 |

**Supplementary Table S6:** Result of GAGE.

| GO | treatment | time (h) | mean_diff | p_val |
| --- | --- | --- | --- | --- |
| GOBP_RESPONSE_TO_INTERLEUKIN_17 | TAA | 24 | 0.550283215 | 1.23085E-07 |
|  |  | 48 | 0.868088354 | 1.71702E-18 |
|  |  | 72 | 0.493649648 | 2.2926E-10 |
|  |  | 96 | 0.366445628 | 4.46203E-06 |
|  |  | 192 | 0.627270248 | 1.57E-07 |
|  | MDA | 24 | 0.861654153 | 7.13E-09 |
|  |  | 48 | 0.964777151 | 5.68E-18 |
|  |  | 72 | 0.775076157 | 1.90E-18 |
|  |  | 96 | 0.565623962 | 3.19171E-11 |
|  |  | 192 | 0.657065653 | 8.34029E-11 |
| GOBP_FIBRINOLYSIS | TAA | 24 | -0.188676844 | 0.024375036 |
|  |  | 48 | -0.61318091 | 4.0677E-05 |
|  |  | 72 | -0.787064263 | 7.15755E-05 |
|  |  | 96 | -0.925128835 | 7.91129E-08 |
|  |  | 192 | -0.206003079 | 7.83472E-08 |
|  | MDA | 24 | 0.69739285 | 2.97E-06 |
|  |  | 48 | 0.58491623 | 2.81E-06 |
|  |  | 72 | 0.59682587 | 6.07E-09 |
|  |  | 96 | -0.003638139 | 0.030206244 |
|  |  | 192 | -0.365541793 | 0.002067302 |

**Supplementary Table S7:** Antibodies used for FACS.

| **marker** | **Detection** | **ID** | **host** |
| --- | --- | --- | --- |
| CD45 | PE | 553081 | Rat |
| CD3 | FITC | 561798 | Rat |
| CD4 | BV786 | 563331 | Rat |
| CD8a | APC-Cy™7 | 561967 | Rat |
| B220 | BV480 | 565631 | Rat |
| NK1.1 | APC | 550627 | Rat |
| $\gamma$ $\delta$ TCR | BV421 | 562892 | Rat |
| CD45 | R718 | 567075 | Rat |
| CD11b | BV480 | 566149 | Rat |
| Ly6C | FITC | 561085 | Rat |
| Ly6G | APC-Cy7 | 560601 | Rat |
| Tim4 | BV421 | 744874 | Rat |
| Siglec-F | Alexa Fluor® 647 | 562680 | Rat |

## Notes

https://github.com/mizuno-group/mda-drinking-water-model

### Summary of Updates

This version has been substantially revised to strengthen the characterization of the MDA drinking-water (MDA-DW) model and clarify its utility for studying cholestatic liver injury. Major updates include additional analyses of drinking-water intake and bile acid-related gene expression, clarification of the experimental design and statistical procedures, and a more detailed discussion of the model's pathological and molecular characteristics. We have also refined the interpretation of the multimodal time-course data, clarified the distinction between cross-sectional and within-animal longitudinal measurements, and revised the manuscript structure to better emphasize the practical utility and limitations of the model. These revisions strengthen the evidence supporting MDA-DW as a complementary model to TAA-DW while preserving the central findings and objectives of the original study.

